# PROS1 released by human lung basal cells upon SARS-CoV-2 infection facilitates epithelial cell repair and limits inflammation

**DOI:** 10.1101/2024.09.11.612489

**Authors:** Theodoros Simakou, Agnieszka M Szemiel, Lucy MacDonald, Karen Kerr, Jack Frew, Marcus Doohan, Katy Diallo, Domenico Somma, Olympia M Hardy, Aziza Elmesmari, Charles McSharry, Thomas D Otto, Arvind H Patel, Mariola Kurowska-Stolarska

**Author notes:** **Corresponding author’s complete name, address, telephone number (including country code, where applicable), and email address** Prof Mariola Kurowska-Stolarska, Sir Graeme Davies Building, 120 University Place Glasgow, G12 8TA. 01413306085.

## Abstract

Factors governing the coagulopathy and pneumonitis associated with severe viral infections remain unresolved. We previously found that the expression of protein S (PROS1) is increased in lung epithelium of patients with mild COVID-19 as compared to severe COVID-19. We hypothesised that PROS1 may exert a local effect that protects the upper airway against severe inflammation by modulating epithelial and myeloid cell responses. To test this, *in vitro* air-interface cultures, seeded from primary healthy human lung epithelial cells, were infected with different SARS-CoV-2 clades. This model, validated by single-cell RNAseq analysis, recapitulated the dynamic cell-profile and pathogenic changes of COVID-19. We showed that PROS1 was located in the basal cells of healthy pseudostratified epithelium. During SARS-Cov-2 infection, PROS1 was released by basal cells, which was partially mediated by interferon. Transcriptome analysis showed that SARS-CoV-2 infection induced proinflammatory phenotypes (CXCL10/11^high^, PTGS2^pos^F3^high^, S100A8/A9^high^) of basal and transitional cells. PROS1 strongly downregulated these cells and transformed the proinflammatory CXCL10/11^high^ basal cells into the regenerative S100A2^pos^KRT^high^ basal cell phenotype. In addition, SARS-CoV-2 infection elevated M-CSF secretion from epithelium, which induced MERTK, a receptor for PROS1, on monocytes added into 3D lung epithelial culture. We demonstrated that SARS-CoV-2 drives monocyte phenotypes expressing coagulation (F13A1) and complement (C1Q) genes. PROS1 significantly downregulated these phenotypes and induced higher expression of MHC class II. Overall, this study demonstrated that the epithelium-derived PROS1 during SARS-CoV-2 infection inhibits the proinflammatory epithelial phenotypes, favours basal cell regeneration, and inhibits myeloid inflammation while enhancing antigen presentation. These findings highlight the importance of basal epithelial cells and PROS1 protection from viral infection induced severe lung pathology.

1) SARS-CoV2 infection of the epithelium results in release of IFN. 2) IFN secretion has an autocrine effect on epithelial cells 3) Infection and IFN cause release of PROS1 from the basal cells, as well as M-CSF from the epithelium 4) PROS1 acts on basal cells which express MERTK, a PROS1 receptor 5) PROS1 downregulated the proinflammatory phenotypes expanded by viral infection, while upregulating KRT^high^ basal cells with repair phenotypes 6) The secreted M-CSF drives MERTK expression on monocytes in cocultures with epithelium. 7) PROS1 induces downregulation of monocyte clusters characteristic of viral infection that express pro-coagulation and complement genes, while upregulating clusters with higher MHC class II. 8) In summary, PROS1 mediates phenotypic switch of SARS-Cov2 induced pathogenic myeloid clusters with complement and coagulation phenotypes into phenotype with efficient antigen presentation, reduces proinflammatory activation of epithelium and induces epithelial barrier repair, resulting in mild COVID-19.

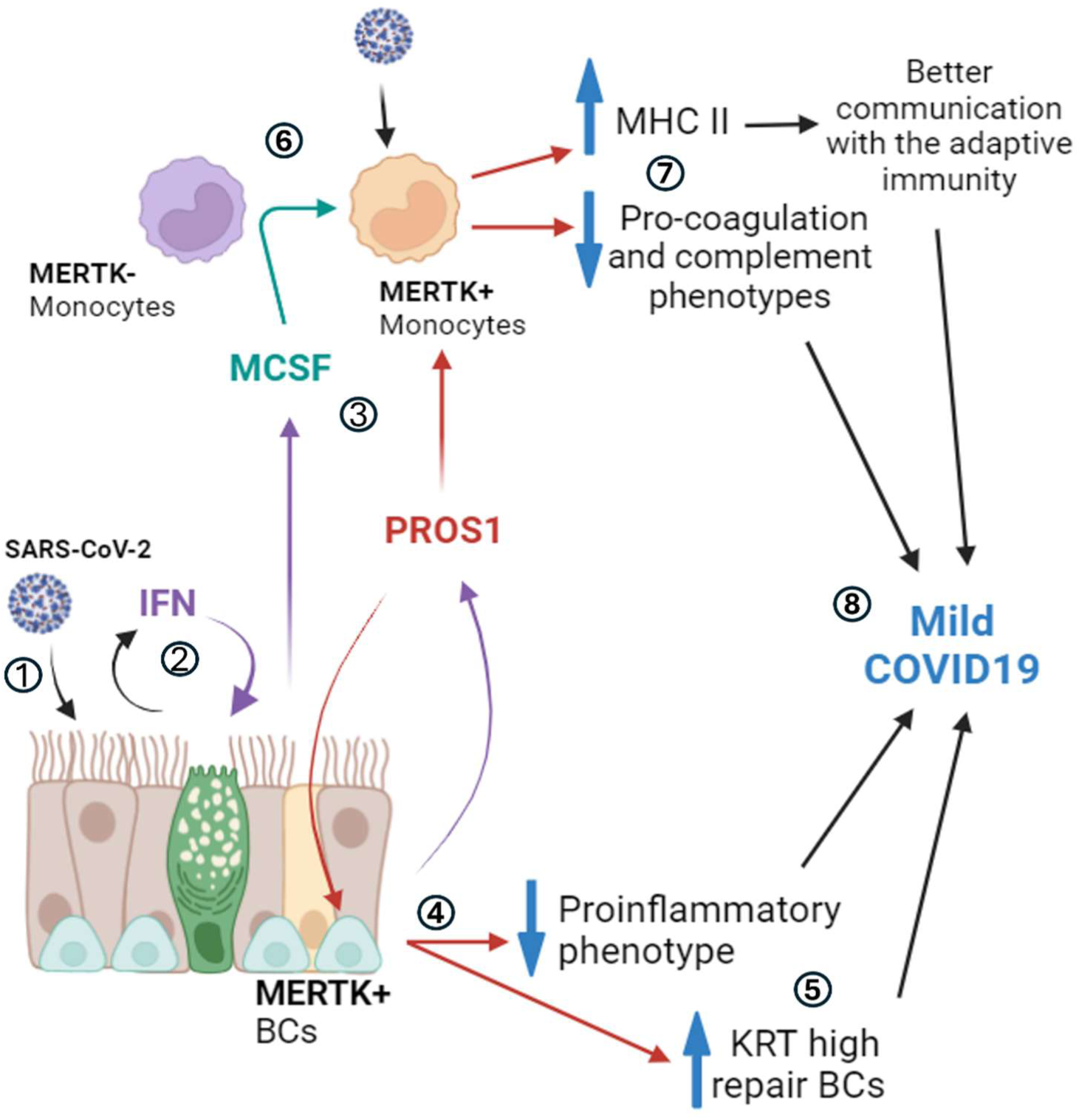

## Introduction

Before the introduction of accessible vaccination against severe acute respiratory syndrome coronavirus 2 (SARS-CoV-2), infection and subsequent COVID-19 illness typically caused 14% of patients to develop severe disease requiring intensive care and oxygen support, while 5% of patients manifested critical disease with life-threatening pneumonia, acute respiratory distress syndrome (ARDS) or septic shock that culminated in multi-organ dysfunction and death (1–3). The severity and mortality of SARS-CoV-2 infection is associated with hyperinflammation and coagulopathy (4), and aberrant activation of myeloid cells in the blood (5–7) and lung (8–10). However, the cellular mechanisms by which disease severity is determined and hyperinflammation and coagulopathy are regulated remain poorly elucidated. COVID-19 remains an ongoing health issue and has resulted in the deaths of 7.06 million people worldwide (WHO, August 2024), so understanding novel cellular mechanisms that ameliorate severe disease in favour of mild symptoms, can help identify pathways that can be targeted for therapy to improve clinical outcomes.

The airway epithelium is an important first line of defence against respiratory viruses, by acting as a physical barrier with tight junctions and mucociliary apparatus, and by releasing multiple antiviral and pro-inflammatory mediators (11–13). The airway epithelial cells are primary portals of entry for SARS-CoV-2 due to expression of ACE2, TMPRSS2, and NRP1 (14–17). They also express multiple pattern recognition receptors, such as the endosomal Toll-like receptors TLR3, TLR7, TLR8 and TLR9, the cytoplasmic receptors retinoic acid-inducible gene I (RIG-I) and melanoma differentiation-associated gene 5 (MDA5), and surface receptors such as TLR2, all of which allow them to detect SARS-CoV-2 and initiate the antiviral interferon response (18,19). Type I and type III IFNs are crucial for the successful defence against SARS-CoV-2 and the prognosis towards mild COVID-19 (12,20–22).

Airway epithelium infected with SARS-CoV-2 interacts with infiltrated monocytes at an early stage of disease and therefore influences disease severity, inducing or modulating overactivation of the myeloid system (23). Mononuclear phagocytes account for 80% of total bronchial alveolar lavage fluid (BALF) cells in patients with severe COVID-19 compared to only 60% in patients with mild disease and 40% in healthy controls (9). Monocytes and macrophages have been shown to play a critical role in the severity of COVID-19 and its features, such as cytokine shock and coagulopathy (4,8).

Myeloid cells interact with the SARS-CoV-2 spike protein using C-type lectins (DC-SIGN, L-SIGN, LSECtin, ASGR1, and CLEC10A) leading to the induction of robust proinflammatory responses that correlate with COVID-19 severity (24). It has also been shown that SARS-CoV-2 can infect and replicate in blood monocytes and lung macrophages (25). About 10% of monocytes and 8% of lung macrophages in patients with COVID-19 get infected with SARS-CoV-2, mostly mediated by CD16 and/or CD64 uptake of opsonized virus (25).

With a focus on epithelial and monocytic involvement in COVID-19, we previously investigated different signatures that could reveal cellular mechanisms influencing disease severity (26). Analysis of single-cell RNA from BALF cells (9) has showed that mRNA encoding the anticoagulant protein *PROS1* was elevated in ciliated epithelial cells in mild, but not severe, COVID-19 (26). Furthermore, analysis of PROS1 in plasma of healthy individuals and patients with mild or severe COVID-19 (with or without treatment) showed no difference (26). This suggested that the PROS1 locally produced in lung airways was protective in COVID-19, and postulates a potential mechanism of immunomodulation by which cytokine shock and severe COVID-19 are prevented.

The vitamin K-dependent anticoagulant Protein S (PROS1) circulates in plasma in two forms: as free protein (30%-40%) and as part of a complex with the complement regulator C4b-binding protein (C4BP) (27,28). Free PROS1 serves as an activated-Protein C cofactor, whereas bound PROS1 can localize C4BP to negatively charged phospholipid membranes (e.g. those exposed on the surface of apoptotic cells), thus providing local control of complement system activation (28).

In addition to its anticoagulation roles, PROS1 is involved in a variety of pathways that affect cell function. PROS1 is an agonist for tyrosine kinase receptors MERTK and TYRO3, through which it regulates immune cell phenotypes, vascular permeability, response to cell damage and coagulation-related pathologies in COVID-19 (29,30). The dysregulation of blood coagulation during COVID-19 results in depleted circulating PROS1 levels (31), which can consequently contribute to cytokine storm by reducing the immunosuppressive action of MERTK in macrophages (32,33) and diminishing intrinsic PROS1 anticoagulant function (28,29,31,34). Subsequently, thromboinflammation can activate the complement system resulting in more inflammation in the lungs of COVID-19 patients (35). SARS-CoV-2 itself can activate the complement system via all three complement pathways (35), and single-nucleotide variants of complement C4BP-α, which interacts with PROS1, are risk factors for morbidity and death in COVID-19 (36).

In airway epithelium the function of PROS1 remain largely unknown. Lung epithelium has been shown to be a local producer of PROS1, upregulating *PROS1* mRNA levels in mild but not severe disease (26). PROS1 has been shown to occupy protective roles in the lung including the prevention of epithelial cell apoptosis and reduction of lung fibrosis (37), which can modulate severity during COVID-19 infection. Similar to observations in COVID-19-associated thrombotic/ischaemic events, SARS-CoV-2 in lung may involve PROS1 cleavage by viral PLpro directly in the site of infection, leading to the local loss of its anticoagulant function (31). Transcriptomic analysis of BALF myeloid cells has demonstrated that infiltrating FCN^+^ and FCN^+^SPP1^+^ monocytes express the PROS1 receptor *MERTK*, suggesting a potential role for PROS1 in modulating myeloid-derived inflammation (9,26).

Therefore, the aims of this study were to: (I) understand the topical production of PROS1 from the bronchial epithelium; (II) elucidate PROS1 roles in modulating epithelial responses during SARS-CoV-2 infection; and (III) study the effect of PROS1 on altering monocyte phenotypes during SARS-CoV-2 infection.

To address these aims, we developed cultures of human primary bronchial epithelial cells grown on air-liquid interface and infected with the SARS-CoV-2 Delta variant. The epithelial cell responses were assessed using single cell transcriptomic analysis, immunofluorescence, and measurement of soluble mediators, including PROS1. We also assessed the expression of MERTK on epithelium and monocytes in cocultures and studied the effect of PROS1 agonism on cell phenotypes.

## Results

### Air-liquid interface bronchial epithelium is capable of replicating SARS-CoV-2 infection and integrates well with tissue biopsies of healthy individuals and patients with COVID-19

Air-liquid interface (ALI) cultures resulted in successful differentiation of pseudostratified bronchial epithelium with cilia and tight junctions (Fig 1, A-D) (Supp Fig 1). We demonstrated that different variants of SARS-CoV-2 (Alpha B.1.1.7, Beta B.1.351, Delta B.1.617.2, Omicron BA.1) could readily infect and replicate in ALI epithelial cells, with viral load increasing progressively from 24 – 72 hours post-infection (Fig 1, E). Since our aim was to study whether PROS1 had ameliorating effects on severe disease, for the Single-cell transcriptomic experiment, we chose the more virulent Delta variant instead of the more recent Omicron variant that causes milder disease (38,39). Based on the transcriptomic results we identified different epithelial cell populations, which we named after their gene expression and upon integration with patient datasets (40) (Fig 1, F-H) (Supp Fig 2, D). All the main epithelial cell phenotypes expected to be found in the upper airway were present in our ALI cultures, including basal, transitional, secretory, deuterosomal and ciliated cells (13,41,42). The culture dataset also integrated well with the published dataset of healthy individuals and COVID-19 patients, validating the *in vitro* system (Fig 1, H, Supp Fig 2).

**Figure 1.**
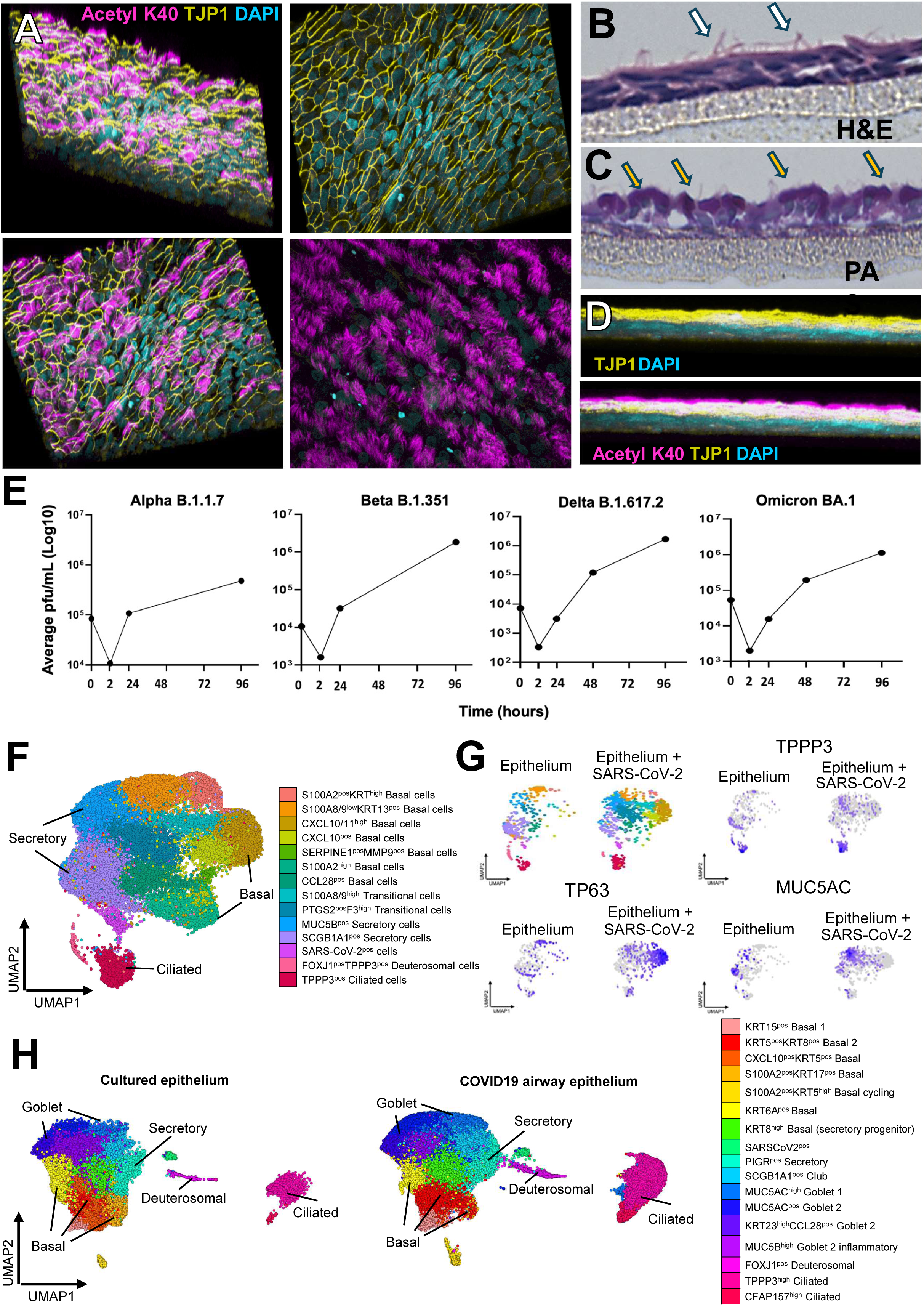
Air-liquid interface cultured epithelium as a system capable of replicating SARS-CoV-2 infection of human bronchial tissues. A. Formation of pseudostratified epithelium with tight junctions (yellow) and cilia (magenta). B. H&E staining of pseudostratified epithelial transverse section with cilia (white arrows); 20X zoomed in. C. PAS staining of the pseudostratified epithelium showing the mucin in red-purple (yellow arrows); 20X zoomed in. D. Lateral view of the 3D section of epithelium from confocal microscopy Z-stacks, showing TJP1 layer (yellow) and the cilia (magenta). E. Infection of the epithelial ALI cultures with different strains of SARS-CoV-2. The 0h represent the virus in the inoculum, the 2h represents the virus in the first wash of epithelium after incubation with the inoculum, and timepoints 24-96h measure the virus that was produced and released from the cells over time. F. UMAP visualization of 36,640 cultured epithelial cells. Each cell is represented by an individual point and is coloured by cluster identity. G. Split UMAP visualization to depict the differences in cluster distribution and TPPP3, TP63 and MUC5AC gene expression between in control vs SARS-Cov-2 infected epithelia. The clusters that were most affected by the SARS-CoV-2 were basal and secretory cells, as shown by the expression of TP63 (basal cell marker) and MUC5AC (secretory cell marker). H. Split UMAP visualization of ALI cultured epithelia (n= 36,640 cells) data integrated with published airway epithelia dataset (n = 63,319 cells) with samples from nasal, tracheal and bronchial epithelium of healthy donors and patients with mild to severe SARS-CoV-2 infection (Yoshida et al 2022).

### PROS1 is expressed and stored in basal cells and is secreted upon SARS-CoV-2 infection

In healthy pseudostratified epithelium, PROS1 is detected in the basal cells by immunofluorescent staining (Fig 2, A). Similarly, in the *in vitro* ALI cultures, PROS1 was expressed in the basal cell layer close to the transwell membrane (Fig 2, B-C). Following infection with SARS-CoV-2 Delta for 72 hours, PROS1 was largely undetected in the basal epithelial cells (Fig 2, C). This was with exception of regions with greater cilia loss, where PROS1 positive cells could be seen aggregated and moving upwards through the pseudostratified tissue (Fig 2, C).

**Figure 2:**
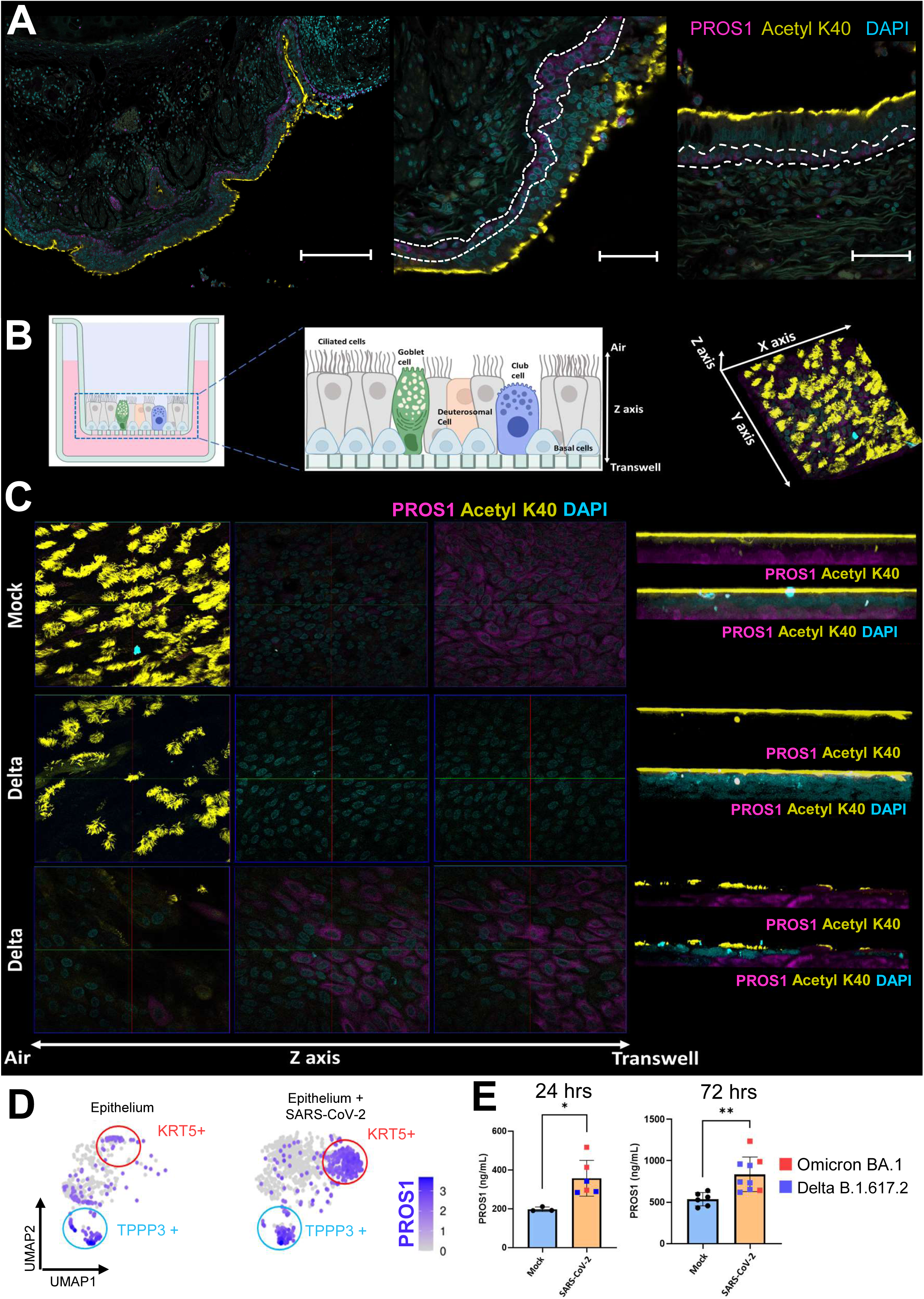
PROS1 is expressed in basal cells of pseudostratified epithelium and secreted upon SARS-CoV-2 infection. A. Basal cells found at the base of pseudostratified epithelium. Tile scan, 20X magnification, scale 250μm. White line indicates basal layer expressing PROS1. B. Stain positive for PROS1 (magenta) in healthy bronchial tissue. Diagram of the Confocal Z scanning of the ALI pseudostratified epithelium. 63X magnification, scale 50 μm. C. PROS1 detection (magenta) in Mock and SARS-CoV-2 (delta strain) infected epithelium across the Z-axis. Left: In Mock (healthy) epithelium PROS1 was detected in the basal cell layer close to the transwell membrane. Infection caused loss of PROS1 from the basal cells. However, in infected epithelia, PROS1 could be detected in areas that had received cilia damage. Right: the lateral view of the 3D scanned tissues corresponding to the ortho-split images on the left. Z stack performed at 63x magnification and analysed using the Zeiss Zen Black software. D. Log-normalised expression of PROS1 visualised on UMAP embeddings of ALI epithelium and ALI epithelium infected with SARS-CoV-2. Highest expression of PROS1 was observed in basal cell clusters (KRT5+, red circle) and ciliated cells (TPPP3+, blue circle). E. Quantification of the secreted PROS1 in mock vs infected epithelium at 24 and 72 hours post infection with SARS-CoV-2. PROS1 was secreted early upon infection from the epithelium and was elevated 72 hours after infection. Each group is represented by multiple transwell systems that were used as control or infected with SARS-CoV-2: Mock 24 hours N= 3, SARS-CoV-2 infected epithelia 24 hours N= 6, Mock 72 hours N= 6, SARS-CoV-2 infected epithelia 72 hours N= 9. Data is presented as bar plot with mean +/- SD. Statistical comparison was performed using unpaired T test between different conditions. *p < 0.05, **p < 0.01, ***p < 0.001, ****p < 0.0001

To determine whether the loss of PROS1 during infection was a result of loss of expression, we interrogated our transcriptomic data and found PROS1 to be expressed in ciliated (TPPP3^pos^) and basal (KRT5^pos^) cells, with the latter being the most affected by the virus infection (Fig 2, D). This indicated that the basal cells still expressed PROS1 mRNA at 72 hours post-infection (Fig 2, D). We next tested whether the cellular loss of PROS1 could be due to its secretion by the basal cells. PROS1 quantification by ELISA in the media of the infected and control ALI cultures was performed at 24 hours and 72 hours post-infection (Fig 2, E). The data showed that infection resulted in increased PROS1 secretion by the infected cells at both timepoints (Fig 2, E), explaining the absence of PROS1 detection in the basal cells by immunofluorescence (Fig 2, C).

### Basal cells express MERTK and TYRO3 and respond to PROS1 by altering proinflammatory phenotypes to pro-regeneration phenotypes during SARS-CoV-2 infection

We next investigated the expression of PROS1 receptors MERTK and TYRO3 in the pseudostratified epithelium (Fig 3, A-B). Immunofluorescent staining of healthy tissue showed expression of MERTK in the basal cells (Fig 3, A), and TYRO3 in all the pseudostratified epithelium (Fig 3, B), indicating that the secreted PROS1 could have autocrine and paracrine effects on the airway cells.

**Figure 3.**
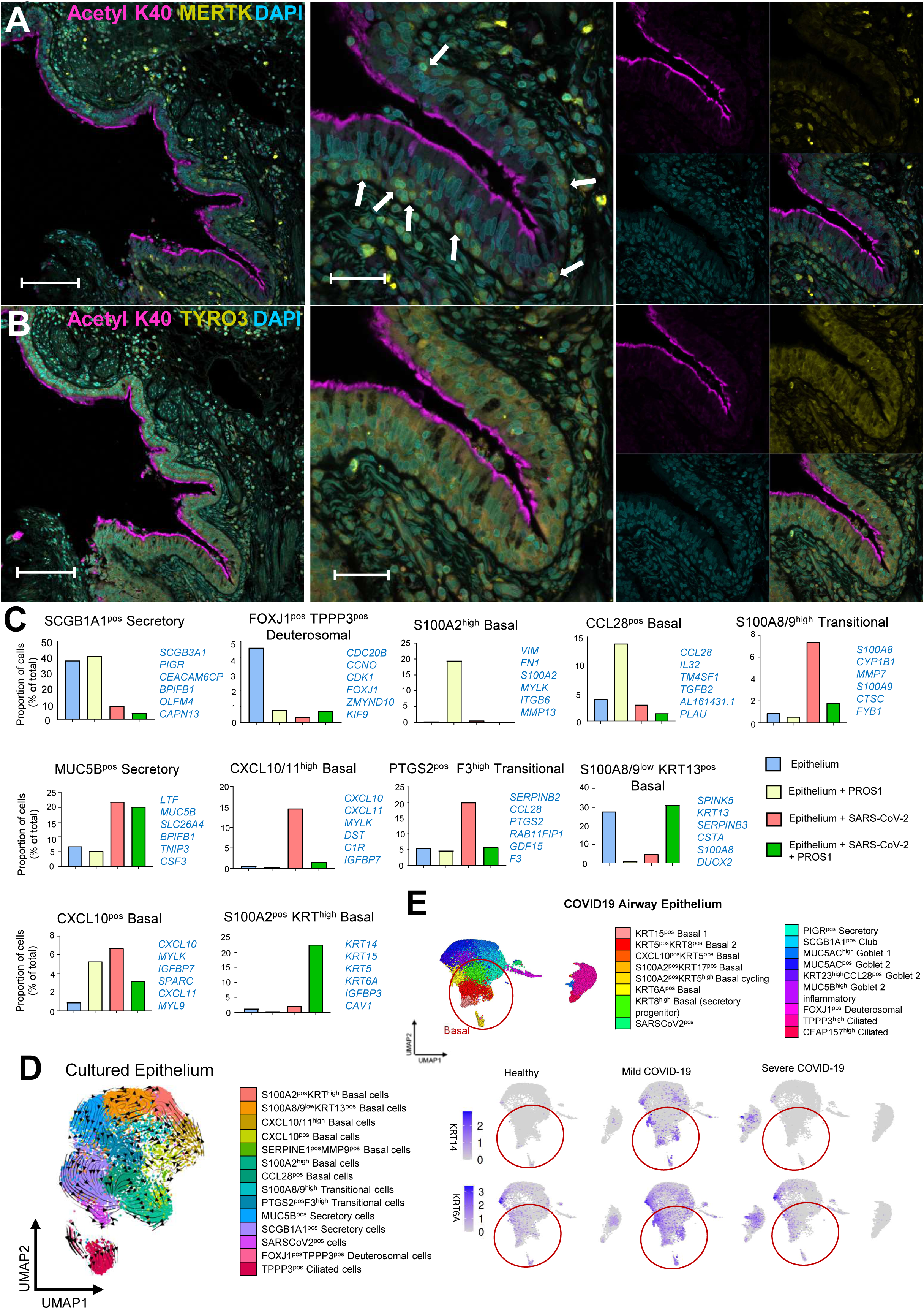
PROS1 acts on basal cells that express MERTK and TYRO1 and ameliorates their proinflammatory phenotype during SARS-CoV-2 infection. A. Left: Expression of MERTK (yellow) in basal cells of healthy pseudostratified epithelium. 20X magnification, scale = 100μm. Middle, Z stack image projection, 63X magnification, scale = 50μm. Right, split channels for Acetylated tubulin (cilia), DAPI (nuclei) and MERTK. Arrows indicate basal cells expressing MERTK. B. Left: Expression of TYRO1 (yellow) in basal cells of healthy pseudostratified epithelium 20X magnification, scale = 100μm. Middle, Z stack image projection, 63X magnification, scale = 50μm. Right, split channels for Acetylated tubulin (cilia), DAPI (nuclei) and MERTK. C. Proportions of cell clusters in control ALI cultures at 72 hours, cultures stimulated with PROS1, cultures infected with SARS-CoV-2, and cultures infected with SARS-CoV2 and treated with PROS1 dissected by scRNAseq, that showed differences between treatments. The proportion of cell clusters per each conditions were obtained by generating a data frame (as.data.frame), using the epithelial object as for idents, and splitting by the SampleID metadata column that represented each condition (Supplemental Table 5). The data were visualised using the png and ggplot functions. The genes listed per each cluster represent top 10 DE gene markers, with a minimum log fold threshold of 0.25. D. Single-cell trajectory of cultured epithelial cells with RNA Velocity analysis visualized on UMAP embeddings. The direction of arrows infers the path of cell trajectory based on spliced versus unspliced RNA counts and suggests a differentiation path from CXCL10/11^high^ basal cells, that were increased during infection, to S100A2^pos^ KRT^high^ basal cells, characterising infected cultures treated with PROS1. E. Integration of ALI scRNAseq dataset with published dataset of COVID19 (Yoshida et al., 2021) showed that KRT14 and KRT6A (markers of PROS1 stimulated epithelium) were increased in basal cells clusters (red circles) in patients with mild COVID-19.

Looking at the proportion of cells we identified different proinflammatory clusters that were characteristic of the SARS-CoV-2 infection, such as the CXCL10/11^high^ basal cells, PTGS2^pos^F3^high^ transitional cells, and S100A8/9^high^ transitional cells (Fig 3, C). Interestingly, PROS1 strongly downregulated the proportions of these phenotypes and favoured the emergence of a S100A2^pos^ KRT^high^ basal cell cluster (Fig 3, C). Characteristic of this cluster was the high expression of cytokeratin genes *KRT14*, *KRT15*, *KRT5* and *KRT6A*. KRT14 and KRT15 are important for the regenerative properties and differentiation of airway basal cells (43). Thus, S100A2^pos^ KRT^high^ basal cells are potentially involved in regeneration of damaged epithelium. When mapping this cluster to the dataset of COVID-19 patients, we observed that this cluster integrated with KRT^high^ cycling cells, which are regenerating cells in lung (Supp Fig 2, C-D).

We then investigated whether the S100A2^pos^KRT^high^ basal cells were an independent phenotype, or whether they were derived from the proinflammatory clusters that were downregulated by PROS1 (Fig 3, C). Trajectory analysis indicated that the S100A2^pos^KRT^high^ cells derived from the CXCL10/11^high^ basal cells (Fig 3, D). This demonstrated that PROS1 transforms the proinflammatory viral-induced basal cell phenotypes to pro-repair and regeneration phenotypes.

We also observed that the expression of *KRT15* and *KRT6A* on the integrated data was higher in epithelial and basal cells derived from patients with mild COVID, compared to those derived from severe COVID-19 (Fig 3, E). This could point to PROS1 favouring mild disease by reducing proinflammatory epithelial phenotypes and by promoting repair of the epithelial barrier.

### Interferon signalling as one of the mechanisms that regulate PROS1 expression and secretion from the bronchial epithelium

After showing that PROS1 is secreted from the epithelium upon infection with the SARS-CoV-2, we wanted to understand the mechanism by which this process occurs.

Knowing that viral infection induces interferon (IFN) response from epithelium, we began by analysing transcription factors downstream of IFN signalling that could bind the PROS1 promoter. ORegAnno analysis (44) showed that STAT1 was a significant regulator of PROS1 by binding its promoter region (Fig 4, A) (Supp table 1). Furthermore, the analysis of regulatory potential of ligands on the epithelial subset from the BALF study (9) predicted that PROS1 could be regulated by multiple upstream ligands, with type 1 and type III interferons revealed to be top genes based on their PROS1 regulatory potential (Fig 4, B).

**Figure 4:**
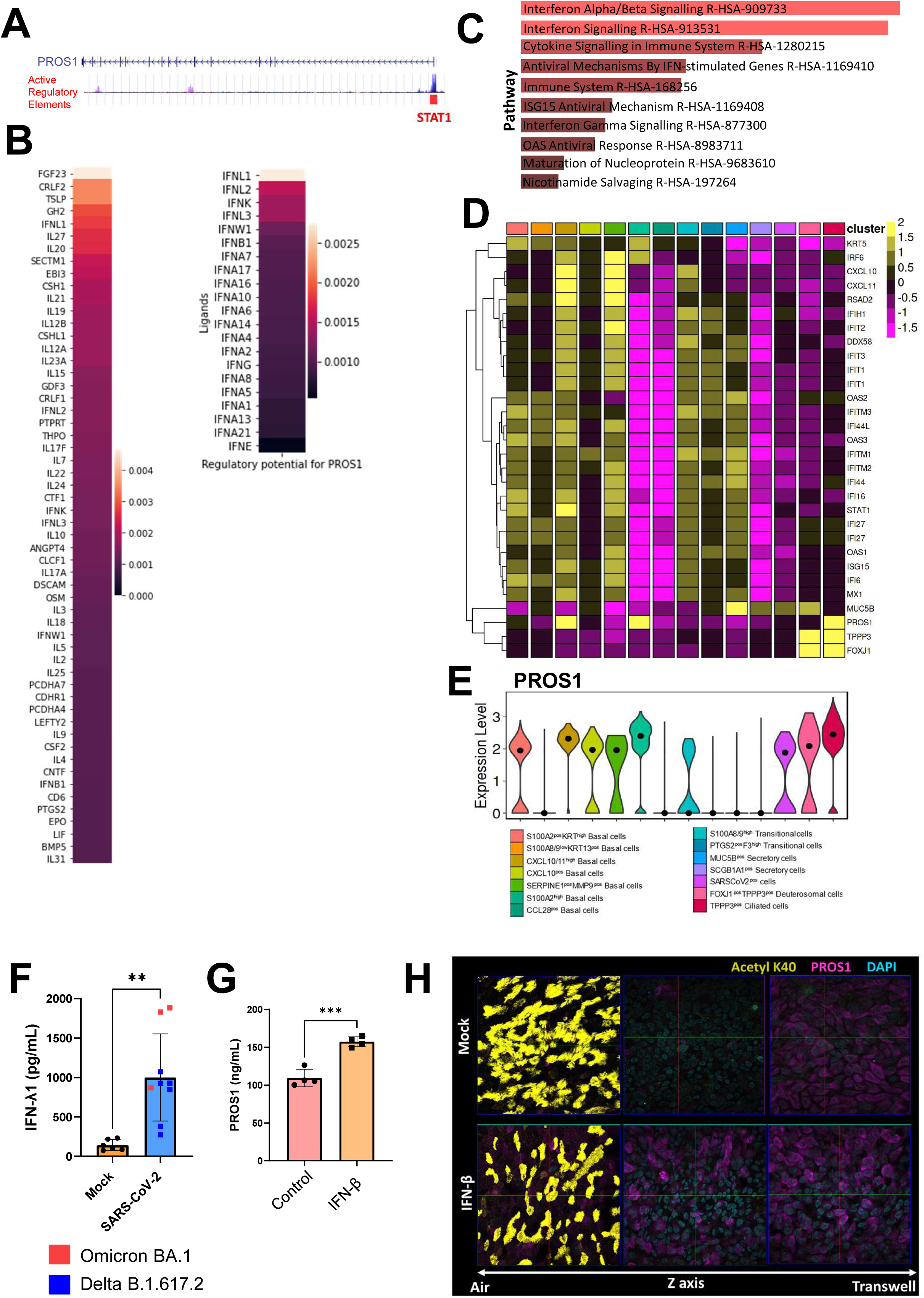
PROS1 is regulated by interferon signalling. A. UCSC Genome Browser visualization of PROS1 gene, the H3K27Ac regions are indicated as active regulatory elements. Multiple transcription factors are reported to bind to PROS1 promoter (ORegAnno annotation), including the STAT1 (indicated) which is involved in interferon signalling and viral responses. Other transcription factors identified included : GATA2, TFAP2C, SMARCA4, CEBPA, TRIM28,CTCF, E2F4, ETS1, FOXA1, FOS, GATA3, HNF4A, TRIM28, MITF, RBL2, SPI1. B. Multiple proteins are predicted to regulate the PROS1 production in epithelial cells. This was done using the ligand-receptor expression analysis (NicheNet), on an object containing only the epithelial cells from the sequenced BALF dataset (Liao et al., 2020). Amongst multiple proteins that could potentially regulate PROS1 expression, either the top regulators (56 ligands with score >0.001) or type I and type III interferons are shown. C. Visualisation of top-10 reactome pathways, ordered by p-value ranking, of epithelial cells infected with virus. Reactome pathway activity inferred by evaluation of differentially expressed genes upregulated in epithelial cells infected with virus vs all other conditions with a minimum log.fc of 0.25 and a p-value < 0.05 based on Wilcoxon Rank Sum Test D. Average-expression heatmap visualising scaled expression of selected DE IFN-inducible genes, alongside putative cluster marker genes, and PROS1 in ALI cultured epithelial cell clusters. DE marker genes for each cluster with a minimum log.fc of 0.25 were identified based on Wilcoxon Rank Sum Test x test with a p-value < 0.05 number (Supp Tab 2). IFN-inducible genes were selected from this list (Supp Tab 2). E. Violin plots illustrating alra-imputed expression values of PROS1 in ALI cultured epithelial cell clusters. Black dot represents median expression value. F. Infection of epithelium with SARS-CoV-2 resulted in Interferon lambda 1 secretion 72 hours after infection. Each group is represented by multiple transwell systems that were used as control or infected with SARS-CoV-2, Mock N= 6, SARS-CoV-2 infected epithelia N= 9. Statistical comparison was performed using unpaired T test between different conditions. *\*p < 0.05, **p < 0.01, ***p < 0.001, ****p < 0.0001*. G. Stimulation of ALI epithelial cultures with 10 ng/mL IFN-β overnight upregulated PROS1 release. Each group is represented by multiple transwell systems that were used as control or stimulated with IFN-β, N= 4. Statistical comparison was performed using unpaired T test between different conditions. *\*p < 0.05, **p < 0.01, ***p < 0.001, ****p < 0.0001*. H. Epithelial ALI cultures stimulated with IFN-β for 24 hours, and then cultured without IFN-β for another 72 hours. The PROS1 (magenta) in IFN-β pre-stimulated cells at 72 hours was present in ciliated cell (Middle Z axis), as compared to the control cultures where it was limited to the basal cells on the surface of the transwell (Left Z axis). PROS1 (magenta), Acetylated tubulin (yellow), DAPI (blue).

Our single cell reactome pathway analyses (45,46) demonstrated that interferon pathways were significantly upregulated in SARS-CoV-2-infected, relative to the mock-infected, ALI cultures (Fig 4, C). Furthermore, our data show that all of the basal cells, which were affected the most during infection, had significantly higher expression of IFN signalling genes and they also had elevated levels of PROS1 mRNA (Fig 4, D). Thus, these results indicate that IFN signalling is a driver of PROS1 mRNA expression in epithelial cells.

We then performed interferon quantification and showed that IFN-λ1 was elevated in the medium of cells at 72 hours post-infection (Fig 4, F), confirming active production upon viral infection that could affect PROS1 secretion (Fig 4, B). To confirm that the IFN could induce PROS1 secretion, we stimulated ALI epithelial cells with IFN-β overnight (Fig 4, G). Though Type III IFN utilizes a receptor complex different from that of type I IFN, both IFN types induce STAT1, STAT2, and STAT3 activation and a similar subset of genes (47). The IFN-β stimulation resulted in higher levels of PROS1 in the media of stimulated cells compared to the controls, indicating IFN signalling induces early secretion of PROS1 (Fig 4, G).

Furthermore, we stimulated epithelial cells for 24 hours with IFN-β to induce activation of IFN pathways, and then cultured them without IFN-β for 72 hours. Using confocal microscopy, we demonstrated that IFN-β pre-stimulation resulted in higher PROS1 in the pseudostratified epithelium and in the ciliated cells, compared to the mock-treated control, where PROS1 was located only in the basal cell layer (Fig 4, H). These observations confirmed that IFN pathway activation results in upregulation of PROS1 production and secretion from the bronchial epithelium.

### Monocytes in coculture with epithelium acquire PROS1 receptor MERTK due to M-CSF production upon viral infection

To investigate the effect of epithelium-derived PROS1 on the monocytes, we performed cocultures of monocytes and ALI epithelial cells (Fig 5 A-C).

**Figure 5:**
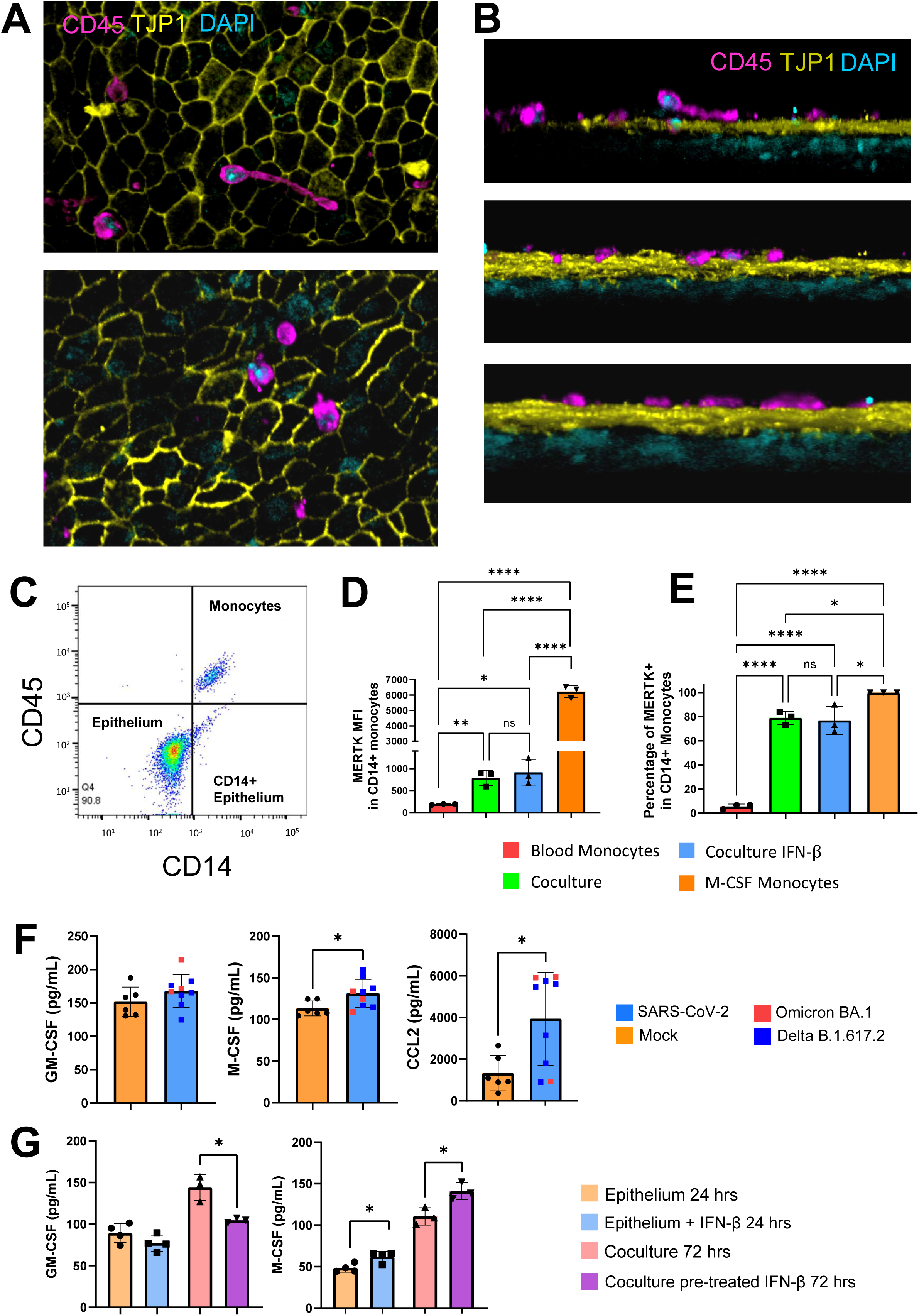
Monocytes interact with epithelium and acquire MERTK in response to M-CSF. A. Confocal 3D scans showing the monocytes on top of the epithelial barrier, at 72 hours cocultures of ALI epithelia with monocytes. B. Lateral view of the 3D scan showing the monocytes on top of epithelial cells, not passing through the tight junctions. C. Dot-plot illustrating that monocytes can be isolated from cocultures with ALI epithelium by expression of CD45 and CD14 D-E. Upon contact with epithelium blood monocytes increased MERTK expression as illustrated MFI of MERTK (D) and by % of MERTK positive cells (E). Each dot represents monocytes from 3 different patients on 3 different ALI epithelial systems (N=3). Statistical analysis was performed using One-Way Anova with Tukey’s multiple comparisons test. *\*p < 0.05, **p < 0.01, ***p < 0.001, ****p < 0.0001.* Data is presented as bar plot +/- SD of mean. F. SARS-Cov-2 Infected bronchial cells produced MCSF and CCL2, but no GM-CSF, at 72 hours. Each group is represented by multiple transwell systems that were used as control or infected with SARS-CoV-2, Mock N= 6, SARS-CoV-2 infected epithelia N= 9. Data is presented as bar plot with +/- SD of mean. Statistical comparison was performed using unpaired T test between different conditions. *\*p < 0.05, **p < 0.01, ***p < 0.001, ****p < 0.0001* G. The IFN-β stimulated epithelium did not affect GM-CSF production. IFN-β pre-treated cocultures had lower levels of GM-CSF compared to controls at 72 hours (left graph). IFN-β stimulation resulted in higher M-CSF production from the epithelium at 24 hours. At 72 hour, the cocultures pre-treated with IFN-β also had higher M-CSF levels than the controls (right). Each group is represented by multiple transwell systems that were used as control or stimulated with 10 ng/mL IFN-β (24 hours stimulated cultures N=4, 72 hours cocultures N= 3). Data is presented as bar plot with mean +/- SD. Statistical comparison was performed using unpaired T test between different conditions. *\*p < 0.05, **p < 0.01, ***p < 0.001, ****p < 0.0001*

We first established that the monocytes expressed the PROS1 main receptor MERTK. Confocal microscopy showed that in the cocultures the monocytes had direct contact with the epithelial cells, migrating and exploring on top of the epithelial barrier (Fig 5 A, B). The monocytes survived well for 3 days in co-cultures and could be readily recovered upon tissue digestion for flow cytometry (Fig 5 C). We then showed that up to 80% of the monocytes co-cultured with epithelial cells for 3 days expressed MERTK (Figs 5D and E). In contrast, the monocytes in circulation did not express MERTK (Fig 5 D). Furthermore, the control M-CSF-treated monocytes expressed MERTK at high levels, with all the cells being positive for the receptor (Fig 5, D, E). In keeping with this, we observed the production of the CSFs from the epithelial cultures infected by SARS-CoV-2. The bronchial epithelium did not produce GM-CSF, but elevated M-CSF at higher levels upon infection than the mock-treated controls (Fig 5, F). The infected epithelia also had higher levels of CCL2, which is the primary chemokine for monocyte recruitment in tissues (Fig 5, F).

We also demonstrated that, similar to infection, M-CSF is higher in cocultures that were pretreated with IFN-β, whereas GM-CSF was largely unaffected by interferon signalling (Fig 5, G). However, the monocytes in cocultures pre-treated with IFN-β did not show a higher level of MERTK (Fig 5 D, E), indicating that M-CSF is one of multiple potential mechanisms that result in MERTK expression.

Collectively, these results demonstrate that SARS-CoV-2 infection causes bronchial epithelium to produce cytokines that invite and support myeloid infiltrates, and induce expression of the PROS1 main receptor MERTK on these cells.

### PROS1 reduces the pro-coagulation and complement-producing monocytes caused by SARS-CoV-2 and favours phenotypes with high expression of THBS1 and MHC class II

Having shown that **SARS-CoV-2** infection causes release of PROS1 from the epithelium, and that the monocytes acquire the PROS1 receptor MERTK in cocultures, we then explored the effects of the virus and PROS1 on the monocyte phenotypes.

Single-cell RNA analysis showed two monocyte phenotypes that were significantly increased proportionally in the presence of SARS-CoV-2 (i.e. F13A1^Pos^ C1Q^High^ and F13A1^Pos^ C1Q^low^ monocytes), and two monocyte clusters that were characteristic of PROS1 stimulation (i.e. HLA^High^ and THBS1^High^ TGFBI^High^ monocytes) (Fig 6 A-C). Interestingly, the virus-induced phenotypes were significantly reduced in the presence of PROS1 (Fig 6, C). In contrast, the PROS1-induced phenotypes were not affected by the presence of the SARS-CoV-2, as they were present in both the cultures supplemented with PROS1 and in cultures supplemented with SARS-CoV-2 and PROS1 (Fig 6 C).

**Figure 6:**
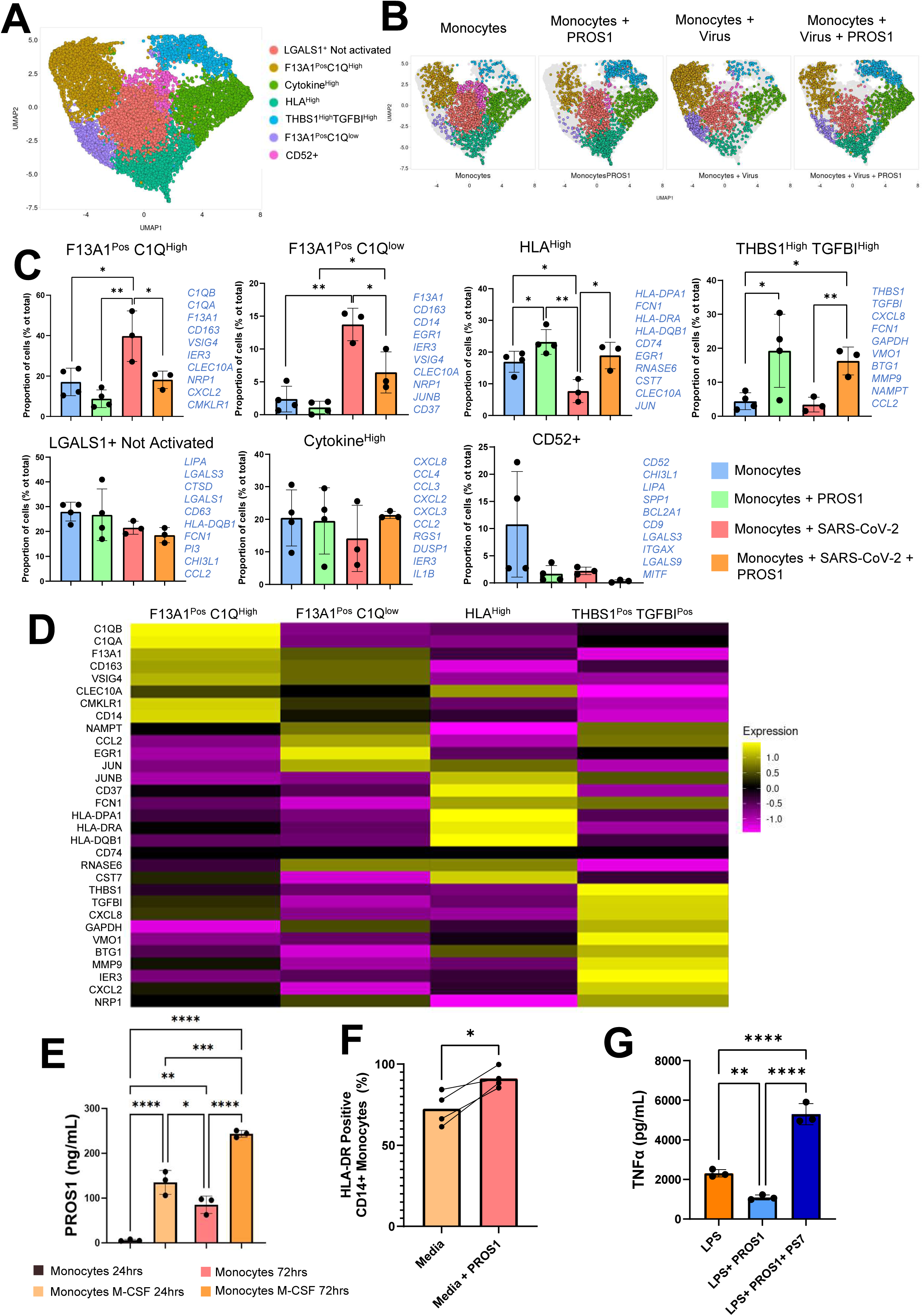
PROS1 downregulates the frequency of monocytes phenotypes with high expression of coagulation and complement genes, and upregulates phenotypes with THBS1 and MHC class II expression. A. UMAP of monocytes cultured for 72 hours with 5 ng/mL M-CSF and in the presence of PROS1 (200 ng/mL), SARS-CoV-2 (10000 pfu), or both. scRNASeq of culture were performed with BD Rhapsody (n=4 different donors). The whole object was normalised and integrated using Seurat SCTransform and the principal component analysis was run. The PCA embeddings were utilised for UMAP generation. The monocytes were subset (9839 cells), the PCA were used to determine the k-nearest neighbours for each cell during SNN graph construction before clustering at the chosen dimension of 22 and resolution of 0.3. B. Split UMAP showing differences in proportion of cell clusters between conditions, as in A). C. Changes in proportion of distinct monocyte phenotypes between multiple conditions is presented as Bar plot with SD of the mean. Proportion of cells clusters as percentage of total cells. Due to small number of cells for D1 (Monocytes+ SARS-CoV-2+ PROS1) and D2 (Monocytes+ SARS-CoV-2) these were removed from analysis, as proportions could not be obtained accurately. For this reason, statistical comparison was performed using unpaired T test between conditions. *\*p < 0.05, **p < 0.01, ***p < 0.001, ****p < 0.0001*. D. Pseudobulk heatmap showing the average expression of DE genes in the four clusters that were affected by PROS1 or SARS-CoV-2. Using the integrated monocyte object (Seurat SCT method), the assay is changed from SCT to RNA. The clusters of interests were subsetted from the rest into a new object. A list of top 10 significant genes (FindAllMarkers, minimum log.fc of 0.25, Wilcoxon Rank Sum Test, adj.p < 0.05) from each cluster was created, including the following genes in order: *C1QB, C1QA, F13A1, CD163, VSIG4, CLEC10A, CMKLR1, CD14, NAMPT, CCL2, EGR1, JUN, JUNB, CD37, FCN1, HLA-DPA1, HLA-DRA, HLA-DQB1, CD74, RNASE6, CST7, THBS1, TGFBI, CXCL8, GAPDH, VMO1, BTG1, MMP9, IER3, CXCL2, NRP1. A*verage expression was performed in the subsetted clusters object (AverageExpression). The heatmap was generated using the object with average expression of genes, and the list of the genes above (DoHeatmap). E. PROS1 (pg/mL) secretion from monocytes without and with M-CSF (50 ng/mL) at 24 and 72 hours of culture. Values are representative of 3 independent experiments (N=3). Statistical analysis performed using One-way Anova with Tukey’s multiple comparison test. *\*p < 0.05, **p < 0.01, ***p < 0.001, ****p < 0.0001,* F. Expression of HLA-DR by CD14 monocytes cultured for 72 hours in media with 5 ng/mL M-CSF and in media with 5 ng/mL M-CSF and PROS1 (200 ng/mL). Values are representative of 4 donors (N=4). Statistical comparison was performed using paired T test. *\*p < 0.05, **p < 0.01, ***p < 0.001, ****p < 0.0001*. G. TNF-α secretion upon overnight stimulation with LPS (1 ng/mL), LPS (1 ng/Ml and PROS1 (200 ng/mL) or LPS 1ng/mL + PROS1 200ng/mL and anti-PROS1 PS7 antibody *10 μg/mL). Macrophages were differentiated for 6 days with 50 ng/mL M-CSF). For stimulation as above, the M-CSF was removed from the media. Data presented as bar-plot of 3 independent experiments (N=3), with SD of the mean. Statistical analysis performed using One-way Anova with Tukey’s multiple comparison test. *\*p < 0.05, **p < 0.01, ***p < 0.001, ****p < 0.0001,*

Furthermore, when analysing the expression of genes averaged per cell clusters using the pseudobulk averaged expression per cell, results showed that PROS1-induced clusters lacked the pro-coagulation and complement genes that were characteristic of the SARS-CoV-2 and had signatures that were not present in both virus-induced clusters (Fig 6, D).

Together, the above results showed that during SARS-CoV-2 infection, PROS1 strongly modulates monocyte phenotypes away from pro-coagulation and complement-producing phenotypes.

We then assessed the endogenous PROS1 production from the monocytes at 24 and 72 hours, in the presence and absence of M-CSF (Fig 6, E). Interestingly, the monocytes cultured without M-CSF did not produce PROS1 at 24 hours cultures but produced it at 72 hours cocultures at levels lower than the M-CSF-cultured cells (Fig 6, E). The cells cultured with M-CSF had PROS1 secretion at both timepoints (Fig 6, E). These results showed that in tissues, monocytes would be exposed to exogenous PROS1 initially, but would also produce PROS1 with time or upon exposure to M-CSF.

To confirm the single-cell mRNA observations on the PROS1-induced phenotypes with high MHC class II expression (Fig 6, C), we analysed monocytes stimulated with PROS1 for 72 hours (Fig 6, F). In accordance with the single-cell mRNA data, PROS1 stimulation of the monocytes resulted in cells with higher MHC class II than the control (Fig 6, F). This indicated that PROS1 upregulated MHC class II at both mRNA and protein level, and could induce monocytes that would communicate better to the effector T cells in tissues.

We also assessed the effects of PROS1 on macrophage activation and TNF-α production. The monocyte-derived macrophages were stimulated overnight with LPS, LPS and PROS1, or LPS and PROS1 and PROS1-inactivating antibody PS7 respectively (Fig 6, G). The results showed that PROS1 reduced TNF-α production from LPS-activated macrophages (Fig 6, G). In contrast, the macrophages produced increased levels of TNF-α in the presence of the PROS1-blocking PS7 antibody (Fig 6, G).

Overall, this data reveals that PROS1 downregulates expression of coagulation and complement genes in monocytes, while favouring the development of cells with elevated expression of MHC class II. We also demonstrated that both exogenous and endogenous PROS1 modulates monocyte-derived macrophage activation in tissues. Control of coagulation and complement, better communication with effector T cells, and regulation of macrophage activation, could be the mechanisms by which PROS1 regulates myeloid inflammatory response.

## Discussion

Cellular mechanisms that promote mild COVID-19 over severe disease have not been elucidated. Our study aimed at investigating the roles of PROS1 in the modulation of COVID-19 severity, by regulating epithelial and monocyte responses during SARS-CoV-2 infection.

Using confocal microscopy and histology of healthy tissues, we showed that PROS1 is located in the basal cells of pseudostratified epithelium (Fig 2 A, C). PROS1 was absent in the SARS-CoV-2 Delta variant-infected cultures, with the exception of cells in areas lacking cilia, where PROS1 was expressed strongly (Fig 2, C). These areas could be most affected by the virus, as SARS-CoV-2 Open Reading Frame 10 (ORF10) protein induces the loss of cilia on epithelial cells by promoting their ubiquitin-dependent degradation (48). Expression of PROS1 in these affected areas may provide protection against rapid disruption of the epithelium, as PROS1 has shown to protect against epithelial apoptosis (37).

We then investigated whether the absence of PROS1 from the basal cells was due to secretion rather than loss of expression. Quantification of PROS1 in the media of infected cultures was higher than in the controls, indicating elevated secretions of PROS1 upon infection (Fig 2, E). This showed that PROS1 was stored in basal cells and slowly secreted in healthy airways, whereas during infection PROS1 was quickly secreted from the basal cells.

In order to assess potential paracrine or autocrine effects of secreted PROS1, we stained for PROS1 receptors MERTK and TYRO3 (29) and found that the healthy tissues’ basal cells express both receptors (Fig 3, A, B). The expression of these receptors could also explain the protective roles that PROS1 exerts on the epithelium (37). Ciliated cells did not express MERTK, but they expressed TYRO3 (Fig 3, A, B). This suggested that the basal cells would be the most affected by PROS1 secretions. In order to dissect the roles of PROS1 on SARS-CoV-2 infected epithelial cells and their phenotypes, we performed Single-cell transcriptomic experiments.

The most affected phenotype from the SARS-CoV-2 were the secretory cells and the basal cell populations (Fig 3, C). An increase in proportion of mucin-producing secretory cells was recorded upon infection, while no difference was observed in the ciliated cells, in accordance with previous published data from SARS-CoV-2 ALI cultures (42). However, we did observe that in the SARS-CoV-2 infected epithelium, unlike in the controls, regions of the pseudostratified epithelium were lacking or having little cilia, as described above (Fig 2,C). This post-translational effect would not necessarily be caught at transcriptome level, or it could also be that the infected ciliated cells were lost upon digestion of tissue.

The viral-induced MUC5B^pos^ secretory cell phenotype, in addition to expressing *MUC5B* that is involved in COVID-19 pathology (49), also expressed genes indicative of viral and proinflammatory signatures. For example, these cells expressed mRNA encoding for *CSF3* and neutrophil chemoattractant *CXCL1, CXCL6, CXCL5* and *CXCL8*, complement system protein *C3, CXCL17* which is a chemoattractant for monocytes and dendritic cells, MHC class I genes (*HLA-A, HLA-C, HLA-E* and *B2M*), and multiple interferon-induced genes (Supplemental Table 2). At protein level, CSF3 (G-CSF) was elevated in infected epithelial cultures at 72 hours (Sup Fig 3, D), whereas CXCL8 (IL-8) was elevated in infected cultures at 24 and 72 hours (Sup Fig 3, J), potentially originating from these secretory cells.

This study also identified three novel phenotypes of basal and transitional cells which were upregulated for SARS-CoV-2 infection, the CXCL10/11^high^ basal cells, PTGS2^pos^F3^high^ transitional cells and the S100A8/A9^high^ transitional cells (Fig 3, C).

The cluster of CXCL10/11^high^ basal cells (KRT5^pos^) were characterised by high expression of *CXCL10* and *CXCL11.* In COVID-19, markedly elevated CXCL10 levels are related to ARDS and neurological complications, such as the loss of smell and taste, and are a good biomarker of disease severity (50–52). CXCL10 and CXCL11 are mostly type II interferon-induced cytokines, however, since our ALI infected epithelial cultures contained no added immune cells, their upregulation could be induced by type I but not type III interferons (53). In fact, multiple interferon inducible genes were upregulated in this cluster (Supplemental Table 2, and Fig 4, D). In addition, this cluster expressed *DDX58* encoding for retinoic acid–inducible gene-I (RIG-I), and *IFIH1* encoding for melanoma differentiation-associated protein 5 (MDA5), that are pattern-recognition receptors for RNA viruses, and downstream induce type I/III interferons (54,55). It has been shown that RIG-I expression levels in human lung cells are important for the cellular defence in the initial SARS-CoV-2 infection, by restraining SARS-CoV-2 replication in a type I/III interferon-independent manner (56). The CXCL10/11^high^ basal cells also had upregulated mRNA encoding for complement components (*C1R* and *C1S*), MHC class I (*HLA-A, HLA-B, HLA-C* and *B2M*) and metalloproteases MMP2 and MMP13, the latter reducing repair during viral infection in bronchial epithelial cells (57).

The PTGS2^pos^F3^high^ transitional cells were characterised by expression of the secretory marker *KRT23* (58), *PTGS2* encoding for cyclooxygenase-2, *PTGES* encoding for prostaglandin E synthase, coagulation factor III (*F3*), and plasminogen activator inhibitor 2 (*SERPINB2*) which may all play roles in COVID-19 inflammatory and coagulation symptoms. This cluster also expresses *EDN1* encoding for endothelin-1, a protein implicated in multiple lung diseases, and in endothelial and cardiovascular dysfunction in severe COVID-19 (59,60) (Supplemental Table 2). Other interesting, upregulated genes of this cluster of cells include the components of tight junction claudin proteins (*CLDN1, CLDN2, CLDN7*), neutrophil chemoattractant (*CXCL1, CXCL5* and *CXCL8*), MHC class I (*HLA-A, HLA-B, HLA-C, HLA-E* and *B2M*), *IFIH1* encoding MDA5 for RNA virus detection, and multiple interferon inducible genes (Supplemental Table 2).

S100A8/A9^high^ transitional cells were characterised by high expression of calprotectin components *S100A8* and *S100A9*. These transitional cells did not express *KRT5* as differentially expressed gene but expressed *KRT6A* and *KRT6B* (Supplemental Table 2), that are stress-induced cytokeratins in airway basal cells with profibrotic phenotypes (Jaeger et al., 2022). These cells expressed other upregulated genes characteristic of SARS-CoV-2 infection that were shared with CXCL10/11^high^ and PTGS2^pos^F3^high^ cells, such as *CXCL11, PTGES, IFIH1*, complement *C1R* and *C1S*, MHC class I encoding genes (*HLA-A, HLA-B, HLA-C, HLA-E, B2M*), metalloproteases *MMP14, MMP7* and *MMP13*, tight junction protein Claudin 1 (*CLDN1*), multiple interferon-inducible genes, and multiple genes encoding ribosomal proteins (Supplemental Table 2).

Interestingly, we observed that these proinflammatory epithelial cells were strongly reduced by PROS1, which favoured a S100A2^pos^KRT^high^ basal cell phenotype. The PROS1 downregulated the virus-induced clusters probably by binding to MERTK and TYRO3, which were shown to be expressed in the basal cells by histology (Fig 3, A, B).

S100A2^pos^KRT^high^ basal cell cluster expanded in the infected cells stimulated with PROS1, but in no other condition. Characteristic of this cluster was the high expression of cytokeratin genes *KRT14, KRT15, KRT5* and *KRT6A*. KRT14 is important for the differentiation of basal cells into secretory and ciliated cells, whereas KRT15 is important for clonogenicity of basal cells (43). Therefore, these two cytokeratins are related to regeneration and repair of airway epithelium (Ievlev et al., 2023). This cluster integrated well with KRT^high^ cycling basal cells from integration with patient dataset (Sup Figure 2, C, D), further supporting their involvement in regeneration of damaged epithelium. We also demonstrated that the KRT14 expression was elevated in basal cells from patients with mild COVID-19 but was not expressed in healthy or severe COVID-19 patients (Fig 3, E). This indicated that the KRT14^pos^ basal cells are important for mild disease phenotype.

We also showed by trajectory analysis that the S100A2^pos^KRT^high^ basal cells originated from the CXCL10/11^high^ basal cells (Fig 3, D), described above as characteristic of viral infection. Multiple genes were upregulated that showed the growth of the S100A2^pos^KRT^high^ cells under infection conditions, such as those encoding interferon-induced proteins, S100 proteins (*S100A2, S100A11, S100A10, S100A8, S100A9, S100A14*) and MHC class I (*HLA-A, HLA-B, HLA-C, B2M*) (Supplemental Table 2). These results showed that PROS1 has the ability to transform the proinflammatory SARS-CoV-2 induced cells to pro-regeneration and pro-repair cells.

Though in our system the PROS1 was artificially added, we demonstrated that PROS1 was secreted by the basal cells upon infection (Fig 2, C, E). We further investigated mechanisms that would results in PROS1 secretion from the epithelium during viral infection. Interferon signalling is a primary antiviral response against SARS-CoV-2 (12,20–22), and the primary signature in our system (Fig 4, C). This was also apparent in the expression of multiple IFN-induced genes in the epithelial cells captured by Single-cell, especially on the basal cells described above (Fig 4, D) (Supplemental Table 2).

Type I IFNs are crucial for the successful defence against SARS-CoV-2 and mild COVID-19, while impairment of their production results in severe disease (20–22). Furthermore, efficient initiation of type III IFN production in the upper airways can lead to rapid elimination of the SARS-CoV-2 and limits viral spread to the lower airways (12). Localisation of IFN response is also important in the lung with regards to disease severity. It has been shown that SARS-CoV-2 induce the efficient production of IFN-III (IFN-λ1 and IFN-λ3) in the upper airways of younger and/or milder patients, whereas critically ill patients express high levels of IFN-I and IFN-λ2 in the lower airways (12). When assessing the IFN production from our infected epithelium, we observed a cumulative secretion of type III IFN, but no type I (Sup Fig 2, G-I). This could be because our epithelial cultures systems were originating from cells of younger individuals, who have shown to produce a type III response against SARS-CoV-2 in the upper airway (12).

We found that STAT1 binds to the promoter region of PROS1, and showed significant regulatory potential based on ORegAnno analysis (44). We further demonstrated IFN-β stimulation of epithelial cells resulted in rapid secretion of PROS1 at 24 hours (Fig 4, G). Furthermore, pretreatment of epithelial cultures with IFN-β overnight to initiate IFN signalling, resulted in elevated PROS1 in the ciliated cells at 72 hours (Fig 4, E). These results together showed that viral induced IFN response is important for the PROS1 production and secretion.

We then observed the effect of secreted PROS1 on the recruited monocytes. The infection of epithelial cultures resulted in elevated M-CSF (Fig 5, F). We demonstrated that CD14+ monocytes acquire MERTK from cocultures with bronchial epithelial cells, due to the M-CSF produced by the epithelium (Fig 5, D, E). This meant that the recruited monocytes, that lack MERTK in circulation (Fig 5, E), when infiltrating the upper airway would be able to respond to PROS1 through MERTK signalling.

SARS-CoV-2 induced two characteristic monocyte clusters with F13A1^pos^C1Q^High^ and F13A1^Pos^C1Q^Low^ phenotypes(Fig 6, C). Both these clusters had expression of *F13A1*, encoding for fibrin-stabilizing coagulation factor 13 alpha chain. Coagulation is a feature of COVID-19, and it has been shown that consumption of the coagulation factor XIIIA (F13A1) was elevated in COVID-19 patients and tended to be higher in non-survivors (61). The two clusters differed at the expression of C1Q genes (Fig 6, D). Complement C1QA and C1QB have shown to be expressed by the monocytes and macrophages in patients suffering from COVID-19 (8,62), and these genes have been associated with complement activation and endothelial dysfunction that is observed in COVID-19 (63).

F13A1^Pos^C1Q^High^ cells also expressed mRNA encoding cytokines and chemokines, such as *CXCL8, CCL2*, and *CXCL2*, showing its roles in recruiting immune cells in lung (Supplemental Table 3). In addition, this cluster expressed *MMP9* (Supplemental Table 3), which impairs epithelial tight junction integrity and reduces tight junction proteins, thus impairing barrier functions (64). In contrast the F13A1^Pos^C1Q^Low^ cells did not produce many cytokines apart from *CCL3*. Both clusters expressed the *NRP1*, encoding for neuropilin1, which potentiates SARS-CoV-2 cell entry and infectivity (14), showing their increased susceptibility for being infected.

Interestingly, PROS1 significantly reduced both these viral-induced clusters when added to the media of cells in presence of SARS-CoV-2, and resulted in two phenotypes of cells, the THBS1^High^TGFBI^High^ and the HLA^high^ monocytes (Fig 6, C).

The THBS1^High^TGFBI^High^ cell cluster had higher expression of mRNA encoding thrombospondin and TGF-β-induced protein. Thrombospondin plays a critical role in efficient wound repair in the lung via the activation of latent TGF-β (reviewed in (65)). The expression of the *TGFBI* on these cells could indicate autocrine TGF-β response, that goes parallel to the THBS1 expression. Thrombospondin also contributes to adequate repair by promoting IL10 production and efferocytosis of apoptotic neutrophils by the macrophages (66). However, the F13A1^Pos^C1Q^High^ and F13A1^Pos^C1Q^Low^ clusters induced by SARS-CoV-2 also expressed THBS1 and TGFBI, though at lower levels (Supplemental Table 3). Furthermore, this cluster also expressed mRNA encoding for multiple proinflammatory mediators such as *CCL2, CCL5, IL1B, CXCL5* and *CCL4*, and *MMP3*. The role of the PROS1-induced THBS1^High^TGFBI^High^ cells remain to be elucidated in the future.

The HLA^High^ cluster, was a cluster of monocytes characterised by the expression of genes encoding for proteins involved in antigen presentation such as MHC class II (*HLA-DPA1, HLA-DRA, HLA-DQB1, CD79*). In addition, this cluster expressed neutrophil recruiting chemokine *CXCL8*, and SARS-CoV-2 entry receptor *NRP1*. The proportion of these cells were elevated for PROS1 supplementation alone and in the presence of SARS-CoV-2 and PROS1, but not in the presence of virus alone (Figure 6, D), showing that the MHC Class II gene upregulation is a strong feature of PROS1 signalling in monocytes regardless of viral infection. Using flow cytometry, we proved that CD14 monocytes cultured with PROS1 upregulated HLA-DR expression on their surface (Fig 6, F). Upregulation of MHC class II can enhance myeloid-T cells communication, allowing for efficient antiviral and effector functions (67,68). T cell responses, such as those including type 1 CD4^+^ T cell phenotype (Th1), are crucial for effective viral control that results in mild COVID-19 (68).

Overall, this study demonstrates that in SARS-CoV-2 infection, PROS1 modulates both epithelial and myeloid responses in the upper airway. In epithelial cells the PROS1 strongly reduces proinflammatory epithelial phenotypes and changes them to pro-regenerative phenotypes that can repair the epithelial barrier. In the recruited monocytes, PROS1 reduces phenotypes with signatures of coagulation and complement, and upregulates

MHC class II phenotypes that may communicate better with antiviral effector T cells. PROS1-induced reduction of epithelial-driven inflammation, promotion of epithelial barrier repair, reduction of myeloid-driven coagulation and complement pathways and efficient antiviral communication between the myeloid and T cells, may all combine to influence mild COVID-19 phenotype. Thus, in this study we describe a novel natural mechanism by which the upper airway regulates inflammation against COVID-19.

## Methods

### Donors and biological variance

The monocytes used in this study were collected from healthy individuals as shown in the table below. To account for sex variance, experiments were designed so that the cells from donors used in experiments came from individuals of different sexes. Furthermore, the donors were selected to be over 45 years of age, as the severe inflammation during COVID-19 was more prominent in older individuals.

| Donor ID | Diagnosis | Sex |
| --- | --- | --- |
| 3458/21 | Healthy | F |
| 3459/21 | Healthy | M |
| 3460/21 | Healthy | F |
| 3461/21 | Healthy | M |
| 3665/21 | Healthy | F |
| 3416/21 | Healthy | M |
| 3542/21 | Healthy | F |
| 3741/22 | Healthy | F |
| 4274/23 | Healthy | F |
| 4290/23 | Healthy | M |
| 4291/23 | Healthy | F |
| 4292/23 | Healthy | M |

### Epithelial Air Liquid Interface (ALI) Culture

Human primary bronchial/tracheal epithelial cells (ATCC, PCS-300-010; Donor: Hispanic/latino man, 14 years old), passage 2, were expanded in T25 flasks in Airway Epithelial Cell Basal Medium (ATCC, PCS-300-030), supplemented with Bronchial Epithelial Cell Growth Kit (ATCC PCS-300-040) and 10 U/mL Penicillin and 10 μg/mL streptomycin (Gibco, 15140-122). After the initial expansion the cells were collected from the flasks and cultured on polycarbonate transwells (NUNC, 140620, 0.4 μm pores, surface area 0.47 cm^2^), at a density of 1×10^5^ cells/cm^2^. Before adding the cells, the transwells were coated with 4.5 μg/cm^2^ Rat tail collagen I (Gibco, A10483-01) in distilled water at 37°C for 6 hours. Following this incubation, the transwells were washed twice with 1X D-PBS (Gibco, 14190-094) and dried at 37°C for another 15 minutes. The cells on transwells were incubated for 4 days, changing the expansion medium after two days.

After the expansion of cells in the transwells, the expansion media was removed, and the apical (cells) and basal compartments of the transwells were washed with warm D-PBS. For the differentiation of epithelial cells in air-liquid interface (ALI) conditions, the Pneumacult ALI differentiation media was used, comprised of Pneumacult ALI Basal Medium (StemCell Technologies, 05002) with 1X ALI Supplement (StemCell Technologies, 05003), and supplemented with 1% ALI maintenance Supplement (StemCell tech 05006), 4 μg/mL Heparin (StemCell Tech. 07980), 0.48 μg/mL Hydrocortisone (StemCell Tech. 07925), and 10 U/mL Penicillin and 10 μg/mL streptomycin (Gibco,15140-122).

The ALI differentiation medium was added to the basal compartment, and cells were cultured at ALI for 4 weeks. The media was changed every two days. When the mucus production was visible, the cells were washed as needed with warm 1X D-PBS. Hydrocortisone was removed from the ALI differentiation medium 3 days before the infection and cocultures. The experiment took place on week 5 of ALI culture.

### Isolation of PBMCs and CD14 monocytes

The PBMCs were isolated from blood of healthy donors using the density separation using Histopaque-1077 (Sigma-Aldrich, 10771-500ML). The blood was diluted 1:2 with sterile 1X D-PBS (Gibco, 14190-094), and then slowly added on top of Histopaque-1077. The blood was centrifuged at 2500 RPM for 25 minutes at room temperature, without break. After the centrifugation, the PBMCs were collected from the buffy coat. The PBMCs were used directly or stored overnight at 4°C before isolation of CD14 monocytes.

CD14 monocytes were isolated using the AutoMACS ProSeparator (Miltenyi Biotech, 130092545) and CD14 labelled beads (Miltenyi Biotech, 130050201), as described in the manufacturer’s protocol. The CD14 positive monocytes were kept on ice in RPMI 1640 media (Gibco, 21875-034), supplemented with 10% FBS (Sigma-Aldrich, F9665), 2mM L-Glutamine (Sigma-Aldrich, G7513-100ML) and 100 U/mL Penicillin and 100 μg/mL streptomycin (Gibco, 15140-122), until ready to be cultured.

### Infection

Epithelial cells cultures were developed as described above and cortisol was removed 3 days before infection. The cells were infected with 10000 pfu of SARS-CoV-2 variants Delta B.1.617.2 and Omicron BA.1 in 100 μL DMEM (DMEM High Glucose GLUTAMax Pyruvate - Gibco 31966021). After 2 hours incubation at 37°C, the inoculum was collected, and the epithelial layers were washed once with 100 μL DMEM (DMEM High Glucose GLUTAMax Pyruvate - Gibco 31966021). The wash was collected as 2 hpi time-point. At 24, 48 and 96 hpi, 100 μL of DMEM was added to the apical layers and incubated at 37°C for 30 minutes. After incubation, the cell washes were stored at −80°C. After all timepoints were collected samples were assayed for virus level by plaque assay on Vero E6 F5 cells, which is a subclone of Vero E6 cells with enhanced susceptibility to SARS-CoV-2 (69).

The same assay was performed earlier, to determine the infectivity of the epithelial cultures towards SARS-CoV-2 Alpha strain B.1.1.7 and Beta strain B.1.351.

### Coculture of epithelium with CD14 monocytes

CD14 monocytes were added on top of epithelium after overnight infection with SARS-CoV-2. The cells were added as a drop of 6 μL containing 1×10^5^ cells on the surface of the epithelial cells. The density of the monocytes relative to the surface area of the epithelium on the transwell was 2.1 ×10^5^ cells/cm^2^. Some of the cocultures were incubated in Pneumacult ALI maintenance medium without hydrocortisone, and some were incubated in the same medium supplemented with PROS1 at 200 ng/mL (Biotechne, 9489-PS). Monocyte-only controls were prepared by seeding 2.1×10^5^ cells/cm^2^ and incubated in complete RPMI medium (5 ng/mL M-CSF) or complete RPMI medium supplemented with PROS1 at 200 ng/mL. The cocultures and controls were incubated at 37°C for 72 hours before single-cell RNA Extraction.

### Culture of monocytes with SARS-CoV-2

To study the direct effect of virus on immune cells, PBMCs from four donors were cultured with SARS-CoV-2 strain BetaCoV/England/02/2020/EPI_ISL_407073 for 72 hours. For this experiment, the cells were seeded at a 2×10^5^ cells/cm^2^ in 24 wells plates and cultured for 72 hours in 400 μL of complete RPMI medium (with 5 ng/mL M-CSF) alone (Control), complete RPMI with 200 ng/mL PROS1, complete RPMI 10000 pfu of SARS-CoV-2, and complete RPMI with10000 pfu of SARS-CoV-2 and 200 ng/mL PROS1. After the incubation period, the cells were collected for RNA extraction from single cells.

### BD Rhapsody Single-cell extraction

On the days of the extraction, the cells were washed with D-PBS, which was collected in individual tubes. Then the cells were lifted using Triple Express (Gibco, 12604-013) for 15 minutes at 37°C. The cell suspension was then transferred into the respective tubes, and the transwells were washed once with D-PBS to collect all the cells. The dead cells were then removed using the EasySep™ Dead Cell Removal (Annexin V) Kit (StemCell Technologies, 17899) as described in manufacturer’s protocol, with the only difference that the cells were collected in 2 mL polypropylene tubes (Sarstedt, 72693) and live cells were negatively separated using DynaMag2 magnet (Life technologies, 12321D), in order to avoid pouring of suspensions while working with the virus.

The cells were then tagged in FACS buffer using the BD Human multiplex tags (BD, 633781). After the incubation, the cells were washed three times, pooled together, and loaded onto the scRNA-seq BD Rhapsody Cartridge of single-cell RNA extraction, using the BD Rhapsody Cartridge Reagent Kit (no. 633731) according to the manufacturer’s protocol. In the final stage, cDNA was synthesised on mRNA captured on the beads using BD Rhapsody cDNA Kit (Cat. No. 633773), following manufacture’s protocol.

### Library formation, mapping and analysis

Libraries of 399 genes were prepared using BD Rhapsody Targeted mRNA and the Tag Amplification Kit (no. 633774) and primers from the BD Rhapsody Immune response Panel Hs (399 genes, BD 633750), as per manufacture’s protocol.

Whole transcriptome libraries were prepared using the Whole transcriptome Analysis and Sample Tag library Preparation kit (BD 633801), as per manufacture’s protocol (2019 version).

Sequencing was performed using Illumina system sequencing services by Glasgow Polyomics.

Mapping was performed in BD Seven Bridges Genomics website. For the target genes, Fastq files containing the raw sequencing were mapped against a reference panel (BD-Rhapsody_immune_Response_Panel_Hs.fasta). For whole transcriptome sequencing, Fastq files containing the raw sequencing were used to map against a reference genome (GRCh38-PhiX-gencodev29-20181205.tar.gz) with supplemental virus genome (Sars_cov_2.ASM985889v3.cds.all.fa), and transcriptome annotation (gencodev29-20181205.gtf).

### Analysis of Epithelium Single cell

Analysis of the data has been performed using the R version 4.1.0. The Seurat package v4 in was used to create an object (CreateSeuratObject funtion) from the RSEC_MolsPerCell files generated from SevenBridges mapping. The data was normalized (NormalizeData) and the top variable genes were identified for all samples (FindVariableFeatures). The data were scaled (ScaleData) and principal component analysis was run (RunPCA).

#### Quality control and filtering

Cell filtering involved removal of cells with less than 200 expressed genes and more than 2000 expressed genes, as well as cells expressing more than 40% mitochondrial genes (subset, subset = nFeature_RNA > 200 & nFeature_RNA <2000 & nCount_RNA > 200 & nCount_RNA < 4000 & percent.mt < 40). This cleaning allowed for exclusion of doublets and dying cells.

#### Clustering pipeline

The PCA embeddings were visualised and then used for UMAP generation (RunUMAP). The same PCs were used to determine the k-nearest neighbours for each cell during SNN graph construction before clustering at the chosen dimension of 26 and resolution of 0.2 (FindNeighbors, FindClusters).

#### Cluster identification and epithelial cluster subsetting

Clusters were identified by their expression of canonical epithelial marker genes (FeaturePlot) and identification of cluster-markers (FindAllMarkers). Epithelial cells clusters were separated into a new object from the co-cultures myeloid cells that were mostly duplets with epithelial cells, and a small cluster of fibroblasts with high expression of collagen genes (subset).

#### Epithelial analysis

The data of the new epithelial object was normalised (NormalizeData) and the top variable genes were identified for all samples (FindVariableFeatures). The data was scaled (Scale Data) and the principal component analysis was run (RunPCA). The PCA embeddings were utilised for UMAP generation (RunUMAP). Clustering pipeline was run as described above, for the chosen dimension of 25 and resolution of 0.5 (FindNeighbors, FindClusters). The clusters and the analysis in the images of this paper are for these clustering. Metadata columns are summarised in the Supplemental Table 5.

Clusters were visualised in UMAP graph (Dimplot) and upregulated genes per each cluster were identified (FindAllMarkers, log fold threshold 0.25). (Supplemental Table 2 with adjusted p-value of <0.05).

#### Enrichr Reactome pathway analysis

Using the epithelial object from above, the idents (Idents) were set to SampleID metadata column, representing different culture conditions such as Epithelial, Epithelial +PROS1, Epithelial + SARS-CoV-2 and Epithelial +SARS + PROS1. The top upregulated genes for each group were obtained (FindAllMarkers, log fold change threshold 0.25) (Supplemental Table 4). The top genes were then used to analyse pathways using the Reactome 2022 in Enrichr website as previously described (45,46,70–72), choosing to display only the statistically significant pathways.

Upstream regulators of PROS1 expression in airway epithelium were predicted NicheNet package (73). Using the following script (https://github.com/saeyslab/nichenetr/blob/master/vignettes/ligand_target_signaling_path.m <u>d</u>), ligand-to-target (PROS1) signalling paths were inferred, and potential ligands were organised by prediction score.

#### Monocyte analysis

SCT Normalization (SCTransform) was used to normalize, scale and find variable features. The samples dataset were integrated using SCT pipeline (SelectIntegrationFeatures, FindIntegrationAnchors), and the principal component analysis was run (RunPCA). The PCA embeddings were utilised for UMAP generation (RunUMAP). Monocytes clusters were separated into a new object (subset). Clustering pipeline was run as described above, for the chosen dimension of 22 and resolution of 0.3 (FindNeighbors, FindClusters). The clusters and the analysis in the images of this paper are for these clustering. Markers were found on SCT assay (PrepSCTFindMarkers, FindAllMarkers with parameters min.pct = 0.4, logfc.threshold = 0.6, recorrect_umi =FALSE).

#### Cell proportion plots

the proportion of cell clusters per each conditions were obtain by generating a data frame (as.data.frame), using the epithelial object as for idents, and splitting by the SampleID metadata column that represented each condition (Supplemental Table 5). The data were visualised using the png and ggplot dev.off functions.

#### Pseudobulk heatmap

Using the integrated monocyte object (Seurat SCT method), the assay is changed from SCT to RNA (DefaultAssay). The clusters of interests were subsetted from the rest into a new object (subset). A list of top 10 significant genes for each cluster is created, listing the genes in order as they would appear on the heatmap. Next, average expression was performed in the subsetted clusters object (AverageExpression). The heatmap was generated using the object with average expression of genes, and the list of the genes above (DoHeatmap).

### Cell Trajectory Analysis

Single-cell trajectory analysis (RNA velocity (74)) of cultured epithelial cells was performed by estimation of spliced and unspliced counts using the velocyto command line interface (velocyto run10x). Outputted .loom files for each culture, which contained transcript splicing data, were incorporated into the epithelial cell Seurat object by splitting the object by culture, loading each .loom file using the (ReadVelocity, SeuratWrappers (0.3.0)) and creating a new assay for spliced, unspliced and ambiguous counts before merging our samples back together again, recreating our integrated Seurat object. This Seurat object was then converted for application in python (SaveH5Seurat, Convert) using SeuratDisk (0.0.0.9019) package. The converted .h5ad file was then read into python using the scanpy (75) (1.9.3) package, which creates an AnnData (0.9.1) object (sc.read). The spliced and unspliced count data was normalized and pre-processed as recommended by scvelo (74) (0.3.1) before running RNA velocity analysis. RNA velocities were estimated for each cell using the (scv.tl.velocity, scvelo), specifying deterministic modelling. Velocities were projected onto the pre-computed UMAP embedding using (scv.pl.velocity_embedding_stream, scvelo).

### Integration with COVID-19 airway epithelium

The generated cultured epithelial dataset was integrated scRNAseq data from nasal, tracheal a nd bronchial airway epithelium of healthy donors and patients with mild to severe SARS-CoV-2 infection (40). The data was obtained in .h5ad format from https://covid19cellatlas.org and was converted using SeuratObject (4.1.3) and SeuratDisk (0.0.0.9015). All paediatric samples were removed along with any cells not of epithelial cell lineage, resulting in dataset of 63,319 cells. Ensemble geneIDs were converted to gene names using gprofiler2 (0.2.1, gconvert). These data w ere then integrated with cultured epithelial data with Harmony () integration. Datasets were mer ged based on expression of 18,525 common genes and the median number of counts per cell between both datasets was identified. This value was used to set scaling factor for combined log-normalization of the merged data (NormalizeData, scale.factor=1926). Feature selection was p erformed (FindVariableFeatures) and the data was scaled (ScaleData) prior to principal compo nent analysis (RunPCA). Cell embeddings from the selected top 40 principal components (PCs) were used in UMAP generation (RunUMAP) to allow for visual inspection of batch separation pri or to integration. Harmony (76) integration was then performed using the Seurat wrapper functi on (RunHarmony, SeuratWrappers, 0.3.0), specifying theta values for sample ID (theta=1) and d ataset (theta=3). The resulting harmony-corrected PCA embeddings were then used for UMAP generation. The same PCs were used for SNN graph construction before clustering at the chosen resolution of 1 (FindNeighbors, FindClusters). Identified clusters were annotated by generating confusion matrix as in MacDonald et al. (77), visualized as a heatmap illustrating the proportion of cells from published airway epithelium reference clusters within each of the new clusters identified post-integration with cultured cells. Differentially expressed genes between clusters wer e identified (FindAllMarkers, test.use=MAST) and such cluster marker genes were identified as those with significant adjusted p-value of <0.05 (Bonferroni and multiple test correction) and expressed by greater than 40% of cells in the cluster (‘min.pct’ parameter 0.4).

### Transwell paraffin embedding

Epithelial cells were fixed in 8% formaldehyde in PBS overnight at RT and stored in D-PBS at 4°C. Before dehydration the D-PBS was carefully removed from the transwell. The fixed cells were dehydrated by incubation with 35%, 50%, 70%, and 95% ethanol for 10 minutes each, and incubation in 100% ethanol twice, 10 minutes each. After the dehydration, the cells were infiltrated twice using Histoclear for 10 minutes each. Infiltration of cells was continued by removing the Histoclear and adding liquid paraffin to the bottom of the well and to the transwell insert. The transwells were incubated at 58°C for one hour. After this incubation, paraffin was changed and transwells were incubated for another hour. After the second incubation, the plates were put at 4°C surface for 20 minutes, to solidify the paraffin. The transwell was carefully removed from the well and the membrane was cut using a small sharp scalper blade. The block containing the transwell membrane was then inserted into a paraffin boat as if it was a piece of tissue, in orientation that would allow transverse sections to be cut. After solidification, the paraffin blocks were kept at RT until cut for the histology.

### Haematoxylin and Eosin staining of transwells paraffin sections

For histological staining, the transwells embedded in paraffin were cut at 5 μm sections. The sections were mounted on 1mm SuperFrost Plus adhesion slides (Epredia, J1800AMNZ). The slides were allowed to dry, and then inserted in an oven for 35 minutes at 60°C. This was followed by dewaxing in Xylene, twice for 3 minutes each. After that the slides were hydrated slowly by incubation in 100%, 90% 70% ethanol, twice and for 3 minutes each. The hydration was completed by incubation in water for 5 minutes.

For H&E staining, the cells were incubated in Harris haematoxylin for 2 minutes and washed with water for 30 seconds. The cells were dipped twice in 1% acid/alcohol solution and rinsed with water. This was followed by incubation in Scott’s Tap Water Substitute for 30 seconds, and a quick rinse in running water. To counterstain, the cells were dipped in 70% ethanol ten times, and then incubated in 1% Eosin for 3 minutes. Following this incubation, the cells were dehydrated using 90% twice for 30 seconds and 100% ethanol twice each for 3 minutes. The cells were incubated in Xylene, twice for 3 minutes each, and then coverslips were mounted using DPX mountant.

### PAS (Periodic Acid Schiff) staining of transwells paraffin sections

The transwell sections were deparaffinize and hydrate to water as described above. The transwells were then oxidize in 0.5% periodic acid solution for 5 minutes. Following a rinse in distilled water, the slides were placed in Schiff reagent for 15 minutes. The slides were then washed in lukewarm tap water for 5 minutes, and counterstained in Mayer’s haematoxylin for 1 minute. The slides were washed in tap water for 5 minutes and dehydrated as described above. The cells were incubated twice in Xylene for 3 minutes each, and then coverslips were mounted using DPX mountant.

### Immunofluorescent staining of ALI cultures on transwells

Virus infected epithelial cells on transwells were fixed in 8% formaldehyde in PBS overnight at RT and stored in D-PBS at 4°C until ready to be stained. The cells fixated with ice-cold 100% methanol and incubated overnight at −20°C. The following day the methanol was removed and transwells were stored dry at 4°C until ready for staining.

The cells were washed 3 times for 3-5 minutes each with D-PBS, and blocked for 1 hours at RT with rocking. Blocking buffer comprised of 1% BSA in 1X D-PBS, 2% Human Serum (Invitrogen, 31876), 2% Goat Serum (Invitrogen, 31872). For permeabilization during blocking, the blocking buffer was supplemented with and 0.2% to 1% Triton X100 for methanol- and PFA-fixated transwells, respectively. The cells were washed 3 times with D-PBS for 3-5 minutes, and the primary antibodies diluted in 1% BSA in D-PBS were added. Antibodies used were: 4 μg/mL rabbit anti-PROS1 (Invitrogen PA5-106880), 2.5 μg/mL rabbit anti-Tight Junction Protein 1 (TJP1) (StemCell Technologies, 100-0750), 1 μg/mL mouse anti-acetylated alpha-Tubulin (StemCell Technologies, 100-0753). For respective isotypes, rabbit IgG (Vector Laboratories, I-1000), and mouse IgG1 (Dako, X0931). The cells were incubated overnight at 4°C with rocking.

After the incubation with the primary antibody, the antibody solution was removed and the transwells were washed 4 times with D-PBS, 5 minutes each. Then the secondary antibodies were added on the transwells diluted in 1X D-PBS at a final concentration of 2 μg/mL. Secondary conjugated antibodies used include anti-rabbit AF647 (Invitrogen, A27040), anti-rabbit AF660 (Invitrogen, A21074), anti-rabbit AF488 (Invitrogen A11008), anti-mouse AF 660 (Invitrogen, A21055) and anti-mouse AF488 (Invitrogen, A28175). The transwells were incubated at RT for 2 hours, in the dark.

After the incubation, the transwells were washed 3X with D-PBS, and then cut into squares using a surgical scalper. The mounting of the transwell section was performed by adding 8 μL of VECTASHIELD DAPI mounting medium (2B Scientific, H-1800-10) on the slide, followed by the transwell membrane with the cells on top. Then another 8 μL of DAPI mounting medium was add on the top, and a coverslip was used to cover the cells. The slides were left to curate initially at RT for 2 hours, and then at 4°C overnight, before observed using confocal microscopy at 63X oil immersion.

For the staining of monocytes in ALI coculture, same technique as described above was used. Antibodies used include mouse anti-human anti-CD45 (2 μg/mL) (Novus-Bio, NBP2-44863), and rabbit anti-human anti-TJP1 (2.5 μg/mL) (StemCell Technologies, 100-0750). For isotype of anti-CD45, mouse IgG2b was used (Novus-Bio, NBP1-43317), whereas for anti-TJP1, rabbit IgG (Vector Laboratories, I-1000) was used. Secondary antibodies used were goat anti-rabbit AF 488 (Invitrogen, A11008) and goat anti-mouse AF660 (Invitrogen, A21055) diluted in 1X D-PBS at a final concentration of 2 μg/mL.

Images were obtained using conformal microscope Zeiss LSM 880, and images acquired using Zeiss Zen Black software.

### Immunofluorescent staining of healthy upper airway tissue

Formalin-fixed paraffin-embedded 5μm-thick upper airway sections were obtained from a healthy biopsy (7123/16). The slides were inserted in an oven for 35 minutes at 60°C. This was followed by dewaxing in Xylene, twice for 5 minutes each. The slides were hydrated by incubation in 100%, 90% 70% ethanol, twice and for 3 minutes each. The hydration was completed by incubation in distilled water for 5 minutes. Epitope retrieval was performed in 0.01M citrate buffer pH 6 and heated using microwave running on 50% power for 5 minutes, and the at 30% power for 8 minutes, until boiling started. The samples were then washed in TBS with 0.025% Triton-X for 5 minutes each and blocked for 2 hours at RT in TBS with 10% human serum, 10% goat serum and 1 % BSA. After blocking, the slides were washed twice in TBS, and primary antibodies or isotopes were added. The slides were incubated overnight at 4°C. The following days, the slides were incubated for 2 hours at RT with the secondary antibodies. Then another 8 μL of DAPI mounting medium was add on the top, and a coverslip was used to cover the cells. The slides were left to curate initially at RT for 2 hours, and then at 4°C overnight, before observed using confocal microscopy at 63X oil immersion.

Antibodies used were: 4 μg/mL rabbit anti-PROS1 (Invitrogen PA5-106880), 1 μg/mL mouse anti-acetylated alpha-Tubulin (StemCell Technologies, 100-0753), 10 μg/mL rabbit anti-MERTK (Abcam, ab52968), 10 μg/mL rabbit anti-TYRO3 (Abcam, ab109231). For respective isotypes, rabbit IgG (Vector Laboratories, I-1000), and mouse IgG1 (Dako, X0931). Secondary conjugated antibodies used at final concentration of 2 μg/mL, as explained above.

### IFN-β Stimulation of ALI Epithelium

For interferon stimulation of the bronchial epithelial cells, hydrocortisone-free ALI differentiation medium was supplemented with 10 ng/mL IFN-β (Peprotech, 300-02BC). The old media was removed, and the IFNB-supplemented media was added in the basal compartment of the transwells. The cells were incubated overnight, and the following day the IFNB-media was removed and collected for ELISA, and transwells washed only in the basal compartment. Then hydrocortisone-free ALI media was added to the transwells and cocultures were set immediately after as described above. As positive controls for MERTK expression, monocytes were seeded in 24 well plates (Corning, 3524), at a density of 1×10^5^ cells/cm^2^, in complete media supplemented with 50 ng/mL human M-CSF (Peprotech, 30025).

### Flow cytometry

On the days of the extraction, the cells were washed with D-PBS, which was collected in individual FACS tubes (Falcon, 352052). Then the cells were lifted using Triple Express (Gibco, 12604-013) for 15 minutes at 37°C. The cell suspension was then transferred into the respective tubes, and the transwells were washed once with D-PBS to collect all the cells. The macrophage controls were also collected using the same method. The cells were then labelled with the viability dye eFluor780 (Invitrogen, 65086514) in D-PBS for 20 minutes in the dark on ice. After the incubation period, cells were washed by adding 1mL of FACS buffer and centrifugation at 1600 RPM for 5 minutes at 4°C. FACS buffer was comprised of 1X D-PBS with 2% FBS (Sigma-Aldrich, F9665), 2mM L-Glutamine (Sigma-Aldrich, G7513-100ML) and 100 U/mL Penicillin and 100 μg/mL streptomycin (Gibco, 15140-122). The supernatant was removed, and cells were labelled with a mix of antibodies against markers of interest for 30 minutes in the dark on ice. The antibodies used were anti-CD14 BV650 (BioLegend, 301835), anti-CD14 AF700 (Biolegend, 301822), anti-CD45 BV711 (BioLegend, 304050), anti HLA-DR (MHCII) BV785 (Biolegend, 307642), anti-MERTK PE-Cy7 (BioLegend, 367610), at 1/100 dilution factor. Unlabelled cells, cells labelled only with eFluor780 (Invitrogen, 65086514) and FMO for each marker of interest were used as technical controls for gating. The data acquisition was performed using BD LSR FORTESSA cytometer. The results were analysed using FlowJo_v10.8.0.

### PROS1 assays on the monocyte-derived macrophages

CD14 monocytes were isolated immediately after PBMCs isolation from whole blood of healthy donors. The PBMCs were isolated as described above. The monocytes were cultured in complete RPMI medium supplemented with 50 ng/mL M-CSF (Peprotech, 30025). The monocytes were differentiated by incubation in M-CSF supplement compete RPMI media for 6 days, changing the medium on day 3. On day six the cells were stimulated with 1 ng/mL LPS (Sigma L6529, clone 055:B5), or with 1ng/mL LPS or 200 ng/mL PROS1 overnight, in absence of M-CSF. Unstimulated and PROS1 stimulated cells were kept as controls. The supernatant for ELISA was collected after the overnight stimulation.

To test the effect of blocking endogenous and exogenous PROS1, we used the anti-PROS1 PS7 antibody (Santa Cruz Biotech, sc-52720) at 10 ug/mL, on macrophages stimulates by LPS (1 ng/mL) or LPS (1ng/mL) and PROS1 (200 ng/mL) overnight.

### ELISA

The media from the lower compartment of the transwells of cocultures and epithelia infected with SARS-CoV-2 was inactivated using a handheld UV Lamp (UVS-28 EI Series UV Lamp, 8 WATT, 254nm, P/N 95-0249-02, 0.32 AMPS, Analytik Jena GmbH). Lamp was placed on top of the plate, and the samples were exposed to UV twice for 2 minutes with a 2 minute break in between exposures. Inactivated medium was collected into centrifuge tubes and stored at −80°C until ready for ELISA. An aliquot of the supernatant was tested in infection assays for the presence of the virus before the supernatant was ready for ELISA.

The media from epithelial cells, cocultures and monocytes treated with IFN-β and the respective controls were collected into centrifuge tubes. The supernatants from cultures where the media was in direct contact with the cells, were centrifuged at 3000 rpm for 10 minutes to ensure no cells were present. After centrifugation, the supernatant was carefully collected into new sterile 1.5mL tubes, avoiding disturbance of the pellet. For transwells, the media from the lower compartment was not centrifuged, because the 0.4 μm pores would not allow cells to pass through the membrane.

Protein S was quantified using the human PROS1/Protein S ELISA Kit (AssayGenie, HUFI01701) as per manufacturer’s protocol.

TNF-α was quantified using human TNFα ELISA kit (Thermofisher, CHC1753), as per manufacturer’s protocol.

CXCL8 was quantified using human IL-8 ELISA kit (Thermofisher, CHC1303), as per manufacturer’s protocol.

The absorbance was measured using a plate reader (BioTek™, ELx800™) and the concentration was calculated using a standard curve generated from the reconstituted standards.

For quantification of multiple cytokines, GeniePlex Multiplex Immunoassays kits were used as instructed by the manufacturers (AssayGenie). The custom human 14-plex consisted of TNF-α, IL-1β, IFN-α2, IFN-β, IFN-λ1(CD29), IL-8 (CXCL8), M-CSF (CSF1), GM-CSF (CSF2), G-CSF (CSF3), Osteopontin (SPP1), Calprotectin (S100A8/A9), IL-6, CCL2 (MCP-1) and IL-10.

### Statistics

Detailed statistical methods are provided in each Figure legend and in the scRNAseq method sections above. For single-cell RNA analysis, the default Wilcoxon Rank Sum Test was used to calculate the DEGs (Seurat, FindAllMarkers).

### Study Approval

This study involved isolation of cells from blood of healthy donors. Written informed consent was received prior to participation.

## Data availability

All scRNAseq data raw files will be available from ArrayExpress under accession number (E-MTAB-14405).

## Author contributions

**TS** Contributed to the study concept, experimental design, generated results, running experiments, writing of the manuscript, optimised workflows, designed protocols, ran libraries for sequencing, developed and ran transwell paraffin embedding protocols, ran histology of paraffin-embedded tissues, performed immunofluorescent staining and transwell preparation for confocal microscopy, ELISA of soluble mediator, Multiplex ELISA, Analysis of single cells data, performed experiments on monocyte cultures and performed multi-parameter flow cytometry.

**AS** Performed the virology experiment, infection of cultures, viral load assays, setting up of cultures, collected samples for analysis, generated results, contributed to writing

**LM** Performed integration with published dataset, analysis of single cells, generated data, contributed to figure and manuscript writing.

**KK** Performed the virology experiments, infection of cultures, viral load assay, collected samples for analysis

**KD** assisted with the flow cytometry and the experiment performance

**DS** contributed to bioinformatic single cell and promoter analysis, and assisted with multiplex ELISA

**JF** Performed trajectory analysis and writing of the manuscript

**MD C**ontributed to the formatting of the images and writing of the manuscript

**OMH** Performed velocity analysis

**AE** Assisted with the immunofluorescent staining of paraffin-embedded lung tissue.

**CM** Contributed to the writing of the manuscript

**TDO** gave access to the servers for bioinformatic analysis

**AHP** initiated the study concept, designed and supervised the overall work, contributed to writing the manuscript

**MKS** initiated the study concept, designed and supervised the overall work, contributed to writing the manuscript.

## Acknowledgments

Funding was supported in part by LifeArc COVID-19 award, and the MRC core award MC_UU_00034/9 (AHP, AS, KK).

