## Supplemental Figures for "PROS1 released by human lung basal cells upon SARS-CoV-2 infection facilitates epithelial cell repair and limits inflammation"

**A**

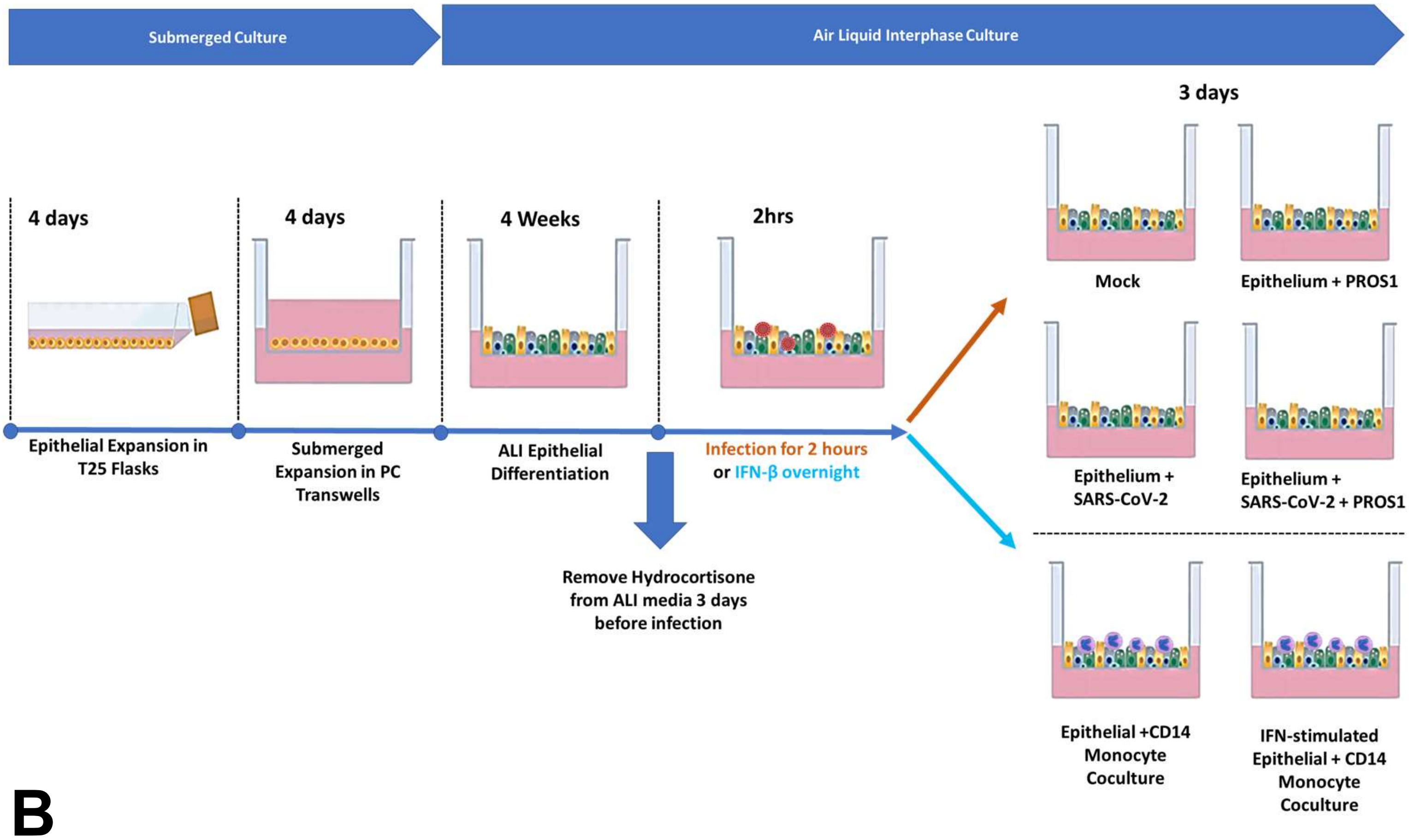
Submerged Culture
Air Liquid Interphase Culture
3 days
4 days
4 days
4 Weeks
2hrs
Mock
Epithelium + PROS1
Epithelial Expansion in T25 Flasks
Submerged Expansion in PC Transwells
ALI Epithelial Differentiation
Infection for 2 hours or IFN-β overnight
Epithelium + SARS-CoV-2
Epithelium + SARS-CoV-2 + PROS1
Remove Hydrocortisone from ALI media 3 days before infection
Epithelial +CD14 Monocyte Coculture
IFN-stimulated Epithelial + CD14 Monocyte Coculture

**B**

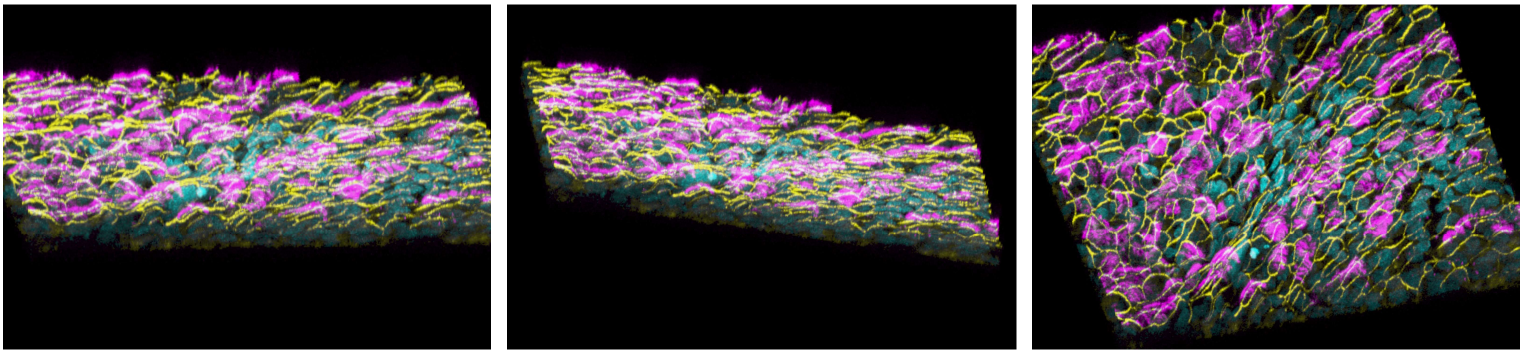

**Supplementary Figure 1: Human Bronchial epithelium grown on Air-Liquid Interface**

A) Methodology of generating pseudostratified epithelium. Human primary bronchial/tracheal epithelial cells (ATCC, PCS-300-010; Donor: Hispanic/latino man, 14 years old), passage 2, were expanded in T25 flasks. After the initial expansion the cells were collected from the flasks and cultured on polycarbonate transwells (0.4 µm pores, surface area 0.47 $cm^2$), at a density of $1x10^5$ cells/$cm^2$. Before adding the cells, the transwells were coated with 4.5 µg/$cm^2$ Rat tail collagen I in distilled water at 37$^o$C for 6 hours. Following this incubation, the transwells were washed twice with 1X D-PBS and dried at 37$^o$C for another 15 minutes. The cells on transwells were incubated for 4 days, changing the expansion medium after two days. After the expansion of cells in the transwells, the expansion media was removed, and the apical (cells) and basal compartments of the transwells were washed with warm D-PBS. For the differentiation of epithelial cells in air-liquid interface (ALI) conditions. The ALI differentiation medium was added to the basal compartment, and cells were cultured at ALI for 4 weeks. The media was changed every two days. When the mucus production was visible, the cells were washed as needed with warm 1X D-PBS. Hydrocortisone was removed from the ALI differentiation medium 3 days before the infection, IFN-β stimulation, and cocultures. The experiment took place on week 5 of ALI culture.

B) At the end of the 4 weeks of ALI culture, the bronchial epithelium was fully differentiated , expressing pseudostratified structure, with tight junctions and cilia. Images were obtained using conformal microscope Zeiss LSM 880, and images acquired using Z stack scanning and analysed using Zeiss Zen Black software.

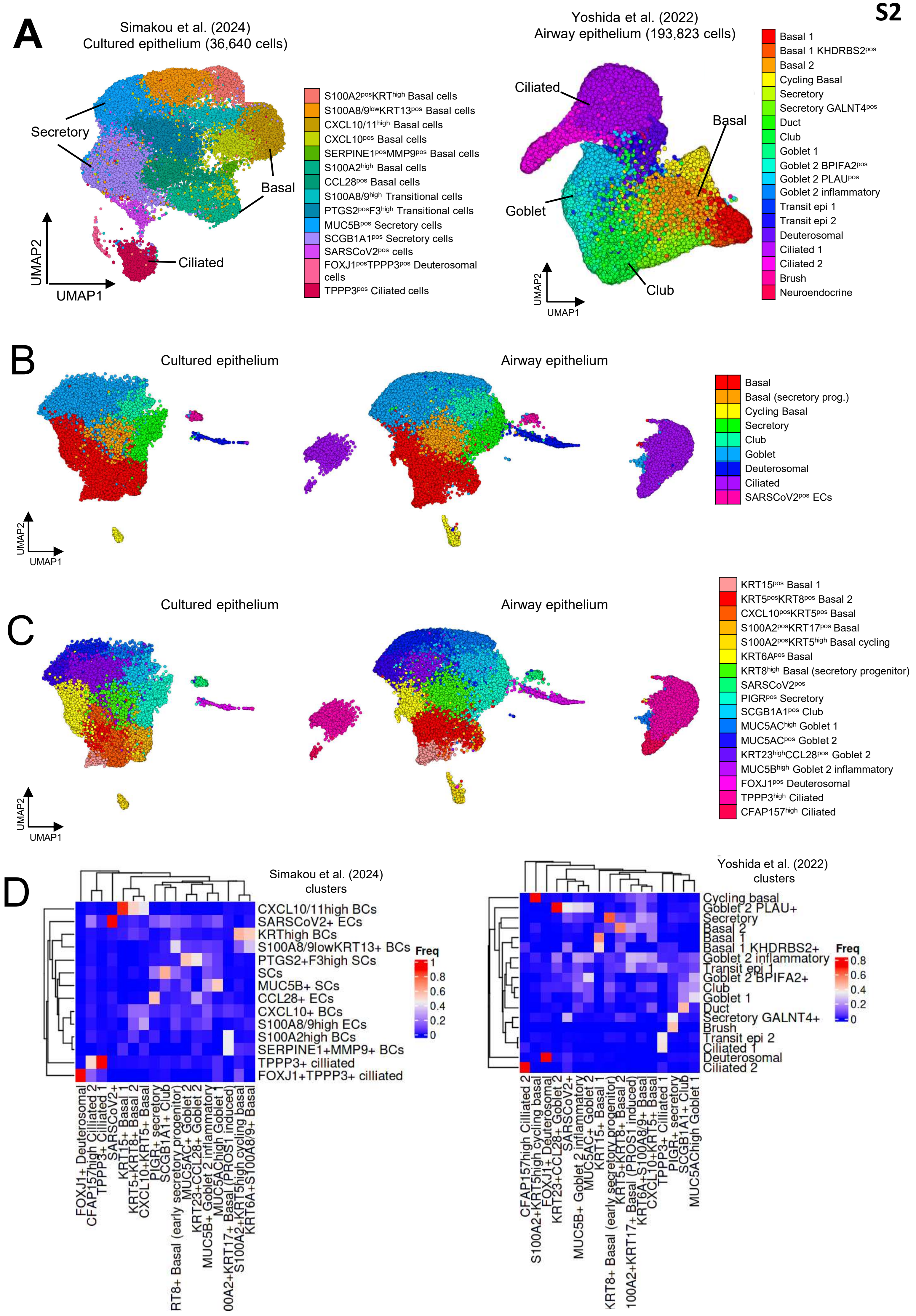
S2
A
Simakou et al. (2024)
Cultured epithelium (36,640 cells)
Secretory
Basal
Ciliated
UMAP2
UMAP1
S100A2posKRThigh Basal cells
S100A8/9lowKRT13pos Basal cells
CXCL10/11high Basal cells
CXCL10pos Basal cells
SERPINE1posMMP9pos Basal cells
S100A2high Basal cells
CCL28pos Basal cells
S100A8/9high Transitional cells
PTGS2posF3high Transitional cells
MUC5Bpos Secretory cells
SCGB1A1pos Secretory cells
SARSCoV2pos cells
FOXJ1posTPPP3pos Deuterosomal cells
TPPP3pos Ciliated cells
Yoshida et al. (2022)
Airway epithelium (193,823 cells)
Ciliated
Basal
Goblet
Club
UMAP2
UMAP1
Basal 1
Basal 1 KHDRBS2pos
Basal 2
Cycling Basal
Secretory
Secretory GALNT4pos
Duct
Club
Goblet 1
Goblet 2 BPIFA2pos
Goblet 2 PLAUpos
Goblet 2 inflammatory
Transit epi 1
Transit epi 2
Deuterosomal
Ciliated 1
Ciliated 2
Brush
Neuroendocrine
B
Cultured epithelium
Airway epithelium
UMAP2
UMAP1
Basal
Basal (secretory prog.)
Cycling Basal
Secretory
Club
Goblet
Deuterosomal
Ciliated
SARSCoV2pos ECs
C
Cultured epithelium
Airway epithelium
UMAP2
UMAP1
KRT15pos Basal 1
KRT5posKRT8pos Basal 2
CXCL10posKRT5pos Basal
S100A2posKRT17pos Basal
S100A2posKRT5high Basal cycling
KRT6Apos Basal
KRT8high Basal (secretory progenitor)
SARSCoV2pos
PIGRpos Secretory
SCGB1A1pos Club
MUC5AChigh Goblet 1
MUC5ACpos Goblet 2
KRT23highCCL28pos Goblet 2
MUC5Bhigh Goblet 2 inflammatory
FOXJ1pos Deuterosomal
TPPP3high Ciliated
CFAP157high Ciliated
D
Simakou et al. (2024) clusters
CXCL10/11high BCs
SARSCoV2+ ECs
KRThigh BCs
S100A8/9lowKRT13+ BCs
PTGS2+F3high SCs
SCs
MUC5B+ SCs
CCL28+ ECs
CXCL10+ BCs
S100A8/9high ECs
S100A2high BCs
SERPINE1+MMP9+ BCs
TPPP3+ cilliated
FOXJ1+TPPP3+ cilliated
Freq
1
0.8
0.6
0.4
0.2
0
FOXJ1+ Deuterosomal
CFAP157high Ciliated 2
TPPP3+ Ciliated 1
SARSCoV2+
KRT15+ Basal 1
KRT5+KRT8+ Basal 2
CXCL10+KRT5+ Basal
PIGR+ secretory
SCGB1A1+ Club
RT8+ Basal (early secretory progenitor)
MUC5AC+ Goblet 2
KRT23+CCL28+ Goblet 2
MUC5B+ Goblet 2 inflammatory
MUC5AChigh Goblet 1
00A2+KRT17+ Basal (PROS1 induced)
S100A2+KRT5high cycling basal
KRT6A+S100A8/9+ Basal
Yoshida et al. (2022) clusters
Cycling basal
Goblet 2 PLAU+
Secretory
Basal 2
Basal 1
Basal 1 KHDRBS2+
Goblet 2 inflammatory
Transit epi 1
Goblet 2 BPIFA2+
Club
Goblet 1
Duct
Secretory GALNT4+
Brush
Transit epi 2
Ciliated 1
Deuterosomal
Ciliated 2
Freq
0.8
0.6
0.4
0.2
0
CFAP157high Ciliated 2
S100A2+KRT5high cycling basal
FOXJ1+ Deuterosomal
KRT23+CCL28+ Goblet 2
SARSCoV2+
MUC5B+ Goblet 2 inflammatory
MUC5AC+ Goblet 2
KRT15+ Basal 1
KRT8+ Basal (early secretory progenitor)
KRT5+KRT8+ Basal 2
100A2+KRT17+ Basal (PROS1 induced)
KRT6A+S100A8/9+ Basal
CXCL10+KRT5+ Basal
TPPP3+ Ciliated 1
PIGR+ secretory
SCGB1A1+ Club
MUC5AChigh Goblet 1

**Supplementary Figure 2:** Integration of the cultured epithelial cells infected with SARS-CoV-2 and stimulated with PROS1 with upper airways of patients with COVID-19

A) UMAP visualization of separate analysis of scRNAseq data from the ALI culture cells (left) and the published dataset (Yoshida et al., 2021) prior to integration.

B) Split UMAP visualization of clusters with coarse cell type annotation of integration of ALI culture cells and published dataset (Yoshida et al., 2021). Both datasets contained the major cells phenotype expected to be present in upper airways, such as the basal, secretory and ciliated cells, and transitional phenotypes (basal to secretory or ciliated) (BéruBé et al., 2010; Mulay et al., 2021; Ruysseveldt et al., 2021)

C) Split UMAP visualization of clusters with fine cell type annotation of integration of ALI culture cells and published dataset (Yoshida et al., 2021).

D) Confusion matrices visualized as heatmaps illustrating the proportion of cells from each cluster ALI culture epithelia scRNAseq (left heatmap) and published airway epithelia reference (right heatmap) clusters within each of the new clusters identified post-integration with Harmony.

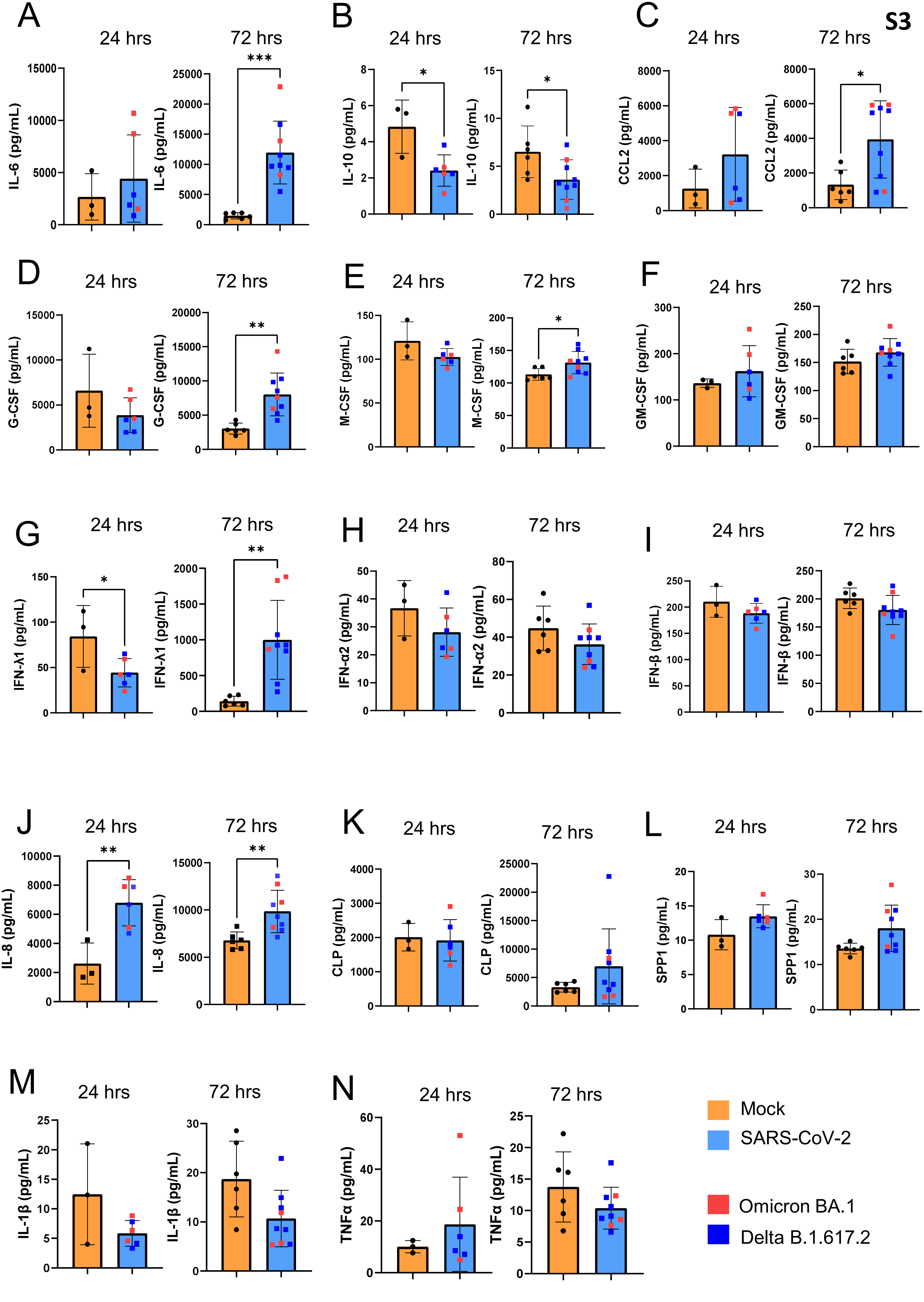
S3
A
B
C
D
E
F
G
H
I
J
K
L
M
N
24 hrs
72 hrs
IL-6 (pg/mL)
IL-10 (pg/mL)
CCL2 (pg/mL)
G-CSF (pg/mL)
M-CSF (pg/mL)
GM-CSF (pg/mL)
IFN-λ1 (pg/mL)
IFN-α2 (pg/mL)
IFN-β (pg/mL)
IL-8 (pg/mL)
CLP (pg/mL)
SPP1 (pg/mL)
IL-1β (pg/mL)
TNFα (pg/mL)
Mock
SARS-CoV-2
Omicron BA.1
Delta B.1.617.2

**Supplementary Figure 3:** Concentrations of proteins of interest in the media of healthy (mock) and SARS-CoV-2 infected epithelia.

A) IL-6 concentrations at 24 hours (left), and 72 hours (right) post-infection.

B) IL-10 concentrations at 24 hours (left), and 72 hours (right) post-infection.

C) CCL2 concentrations at 24 hours (left), and 72 hours (right) post-infection.

D) G-CSF concentrations at 24 hours (left), and 72 hours (right) post-infection.

E) M-CSF concentrations at 24 hours (left), and 72 hours (right) post-infection.

F) GM-CSF concentrations at 24 hours (left), and 72 hours (right) post-infection.

G) IFN-λ1 (IL-29) concentrations at 24 hours (left), and 72 hours (right) post-infection.

H) IFN-α2 concentrations at 24 hours (left), and 72 hours (right) post-infection.

I) IFN-β concentrations at 24 hours (left), and 72 hours (right) post-infection.

J) IL-8 (CXCL8) concentrations at 24 hours (left), and 72 hours (right) post-infection.

K) Calprotectin (S100A8/A9) concentrations at 24 hours (left), and 72 hours (right) post-infection.

L) SPP1 concentrations at 24 hours (left), and 72 hours (right) post-infection.

M) IL-1β concentrations at 24 hours (left), and 72 hours (right) post-infection.

N) TNF-α concentrations at 24 hours (left), and 72 hours (right) post-infection

Each group is represented by multiple transwell systems that were used as control or infected with SARS-CoV-2, Mock N= 6, SARS-CoV-2 infected epithelia N= 9. Data shown as bar-plots with SD of the mean. Statistical analysis between the condition performed using unpaired T test, and p values < 0.05 were considered significant. *$^*p < 0.05$, $^{**}p < 0.01$, $^{***}p < 0.001$, $^{****}p < 0.0001$*
